# Distinct *MUNC* lncRNA structural domains regulate transcription of different promyogenic factors

**DOI:** 10.1101/2021.06.22.449443

**Authors:** Roza K. Przanowska, Chase A. Weidmann, Shekhar Saha, Magdalena A. Cichewicz, Kate N. Jensen, Piotr Przanowski, Patrick S. Irving, Michael J. Guertin, Kevin M. Weeks, Anindya Dutta

## Abstract

Many lncRNAs have been discovered using transcriptomic data, however, it is broadly unclear what fraction of lncRNAs are functional and what structural properties affect their phenotype. *MUNC* lncRNA, also known as ^DRR^eRNA, stimulates skeletal muscle differentiation. The prevailing hypothesis is that *MUNC* stimulates the *Myod1* gene *in cis* as an enhancer RNA and stimulates expression of several other promyogenic genes *in trans* by recruiting the cohesin complex to their promoters. Experimental probing of the RNA structure revealed that *MUNC* contains multiple structural domains not detected by RNA structure prediction algorithms in the absence of experimental information. We discovered that these specific and structurally distinct domains are required for induction of different promyogenic genes, for binding at different genomic sites to regulate the expression of adjacent genes, and for binding the cohesin complex. Moreover, we found that induction of *Myod1* or interaction with cohesin comprise only a subset of the broad regulatory impact of this lncRNA. Our study thus reveals unexpectedly complex, structure-driven functions for the *MUNC* lncRNA and emphasizes the importance of experimentally determined structures for understanding structure-function relationships in lncRNAs.

## Introduction

In 2004 the International Human Genome Sequencing Consortium found that roughly 70% of the nucleotides in the human genome are transcribed but not translated, thus comprising non-coding RNAs (ncRNAs) (Consortium, 2004). Since then, many have been discovered and characterized, and a few of them functionally characterized, but even fewer have experimentally determined structural information. The majority of long non-coding RNAs (lncRNAs), defined as transcripts longer than 200 nucleotides, share most signatures of coding RNAs: they are transcribed by RNA polymerase II, spliced, capped at the 5′ end with 7-methylguanosine, and poly-adenylated at the 3’ end (Derrien et al., 2012). lncRNAs are not transient intermediaries to functional proteins, but functional macromolecules that drive cellular programs, therefore it is critically important to understand how the intrinsic lncRNA structure enables higher-order complex formation to fulfil its function.

lncRNAs can activate (Wang et al., 2014) or repress (Kino et al., 2010) gene expression. Enhancer RNAs (eRNAs) are a subclass of lncRNAs that facilitate chromatin looping and transcription, mostly *in cis* (Lai et al., 2013; Ørom et al., 2010). lncRNAs can mediate epigenetic changes by directly recruiting chromatin remodeling complexes to specific genomic loci (Mondal et al., 2015; Postepska-Igielska et al., 2015; Rinn et al., 2007). lncRNAs are frequently reported to act as sponges that compete for binding to miRNAs and proteins, consequently blocking mechanisms of post-transcriptional control (Elguindy et al., 2019; Munschauer et al., 2018; Szcześniak and Makałowska, 2016). lncRNAs can create subcellular domains, scaffolding the assembly of small ribonucleoprotein complexes (Tsai et al., 2010; Yoon et al., 2013) and larger nuclear bodies (Chujo et al., 2016; Yamazaki and Hirose, 2015). Some lncRNAs have established roles in disease, for example, acting as tumor suppressors (Sakurai et al., 2015) or tumor promoters (Reon et al., 2018).

Despite the fact that lncRNAs have been studied for over 30 years, little is known about structures of lncRNAs or about how these structures contribute to function. Multiple technologies have been employed to model structures of lncRNAs. These methods fall into three major categories: bioinformatic modeling, biochemical probing, and high-resolution methods. Most published studies have predicted the secondary structures of lncRNAs computationally by leveraging thermodynamic properties of base pairing, base stacking, and other atomic interactions. Although modeling can be quite accurate for short RNA sequences, accuracy drops significantly as the length of the transcript increases (Deigan et al., 2009; Miao et al., 2015). Biochemical probing technologies rely on small-molecule or enzyme probes that react selectively with structured or unstructured RNA. Data from probing experiments can be used to restrain folding algorithms to yield more accurate structural models than result from computational prediction alone, especially for long RNAs (Li et al., 2020a). SHAPE-MaP chemical probing strategies have proven especially useful as nearly every nucleotide is probed in a single experiment and RNAs of nearly any length can be studied (Busan et al., 2019; Merino et al., 2005; Smola et al., 2015a; Smola et al., 2015b). Chemical probing studies have revealed that individual RNAs have “structural personalities” (Weeks, 2021), a feature that likely applies to lncRNAs, and to individual domains within large lncRNAs. Structure probing of lncRNAs and connecting defined motifs to a defined function is in its infancy. High-resolution methods, especially of ribosome (Watson et al., 2020) and viral (Jaafar and Kieft, 2019) systems have revealed enormous complexity in RNA structure and that large RNAs tend to form smaller domains.

Classification and characterization of novel lncRNAs usually starts by establishing their subcellular localization, stability, specificity, and abundance, followed by examining their function. Similar to proteins, RNA is expected to exert its biological function, in part, through specific structure. Lack of the structure-function studies in the lncRNA field is a major limitation for determining the molecular mechanisms by which lncRNAs exert their functions. One of the best-characterized lncRNAs is *Xist* lncRNA, which is a master regulator of X chromosome inactivation (Brown et al., 1991; Cerase et al., 2015). Deletion studies showed that a 5’ conserved repeat region (RepA) of *Xist* is indispensable for gene silencing (Wutz et al., 2002). NMR studies found that a 26-nucleotide fragment that includes the RepA sequence forms a stem-loop structure (Duszczyk et al., 2008). More recent studies have further demonstrated the influence of structure on the functions of lncRNAs: Dynamic and flexible structures in *Xist* act as landing pads for proteins (Fang et al., 2015) (Smola et al., 2016), *MEG3* pseudoknot structures (“kissing loops”) modulate the p53 response (Uroda et al., 2019), an unstructured region in the *SLNCR1* lncRNA nucleates a non-canonical transcription complex that promotes melanoma invasion (Schmidt et al., 2020), and *GAS5* lncRNA contains three structural domains that independently regulate cell survival under different conditions (Frank et al., 2020).

The *MUNC* lncRNA (also known as ^DRR^eRNA) plays an important role in myogenesis, the process of skeletal muscle tissue formation (Cichewicz et al., 2018; Mousavi et al., 2013; Mueller et al., 2015; Tsai et al., 2018). *MUNC* is located 5-kb upstream of the *Myod1* transcription start site and has two functional isoforms (Mueller *et al*., 2015). Both are upregulated in skeletal muscles compared to other tissues and during differentiation of skeletal muscle myoblasts (Mueller *et al*., 2015). *MUNC* was initially thought to be a classic eRNA that acts to maintain open chromatin and induce expression of the *Myod1* gene *in cis*. However, *MUNC* depletion using small interfering RNA (siRNA) reduces *Myod1* transcription and myoblast differentiation, and since siRNA acts post-transcriptionally, this effect is inconsistent with a *cis*-acting effect on transcription. Consistent with this *cis* activity, stable overexpression of *MUNC* lncRNA from heterologous loci stimulates the expression of multiple promyogenic RNAs, including *Myod1*, meaning that the *MUNC* lncRNA operates *in trans* (Mueller *et al*., 2015). *MUNC* overexpression in C2C12 cells that lack the *Myod1* gene and therefore the MYOD1 protein induces the expression of other myogenic genes, demonstrating that *MUNC* is capable of regulating genes in a MYOD1-independent manner (Cichewicz *et al*., 2018). Thus, *MUNC* regulates gene expression *in cis* and *in trans*. It has been suggested that the *trans* functions of *MUNC* are mediated through the recruitment of the cohesin complex to target promoters (Tsai *et al*., 2018). Here, we describe the first structure-function study of *MUNC* lncRNA. We characterized multiple RNA functional domains in *MUNC* and found that different domains mediate distinct features of *MUNC* promyogenic activity.

## Results

### MUNC lncRNA Promotes Promyogenic Pathway Differentiation

The *MUNC* gene encodes two functional lncRNA isoforms that are approximately equal in abundance (Mueller *et al*., 2015), that differ by the inclusion of an intron, which we refer to as spliced and unspliced. We stably overexpressed each isoform separately in C2C12 murine myoblasts and cultured the resulting cell lines in proliferating (GM) or differentiating (DM3) conditions for 3 days (Figure 1A). Each isoform induced expression of *Myod1*, *Myog*, and *Myh3,* though expression of the spliced isoform resulted in higher levels of these transcripts (Figure 1B).

**Figure 1.**
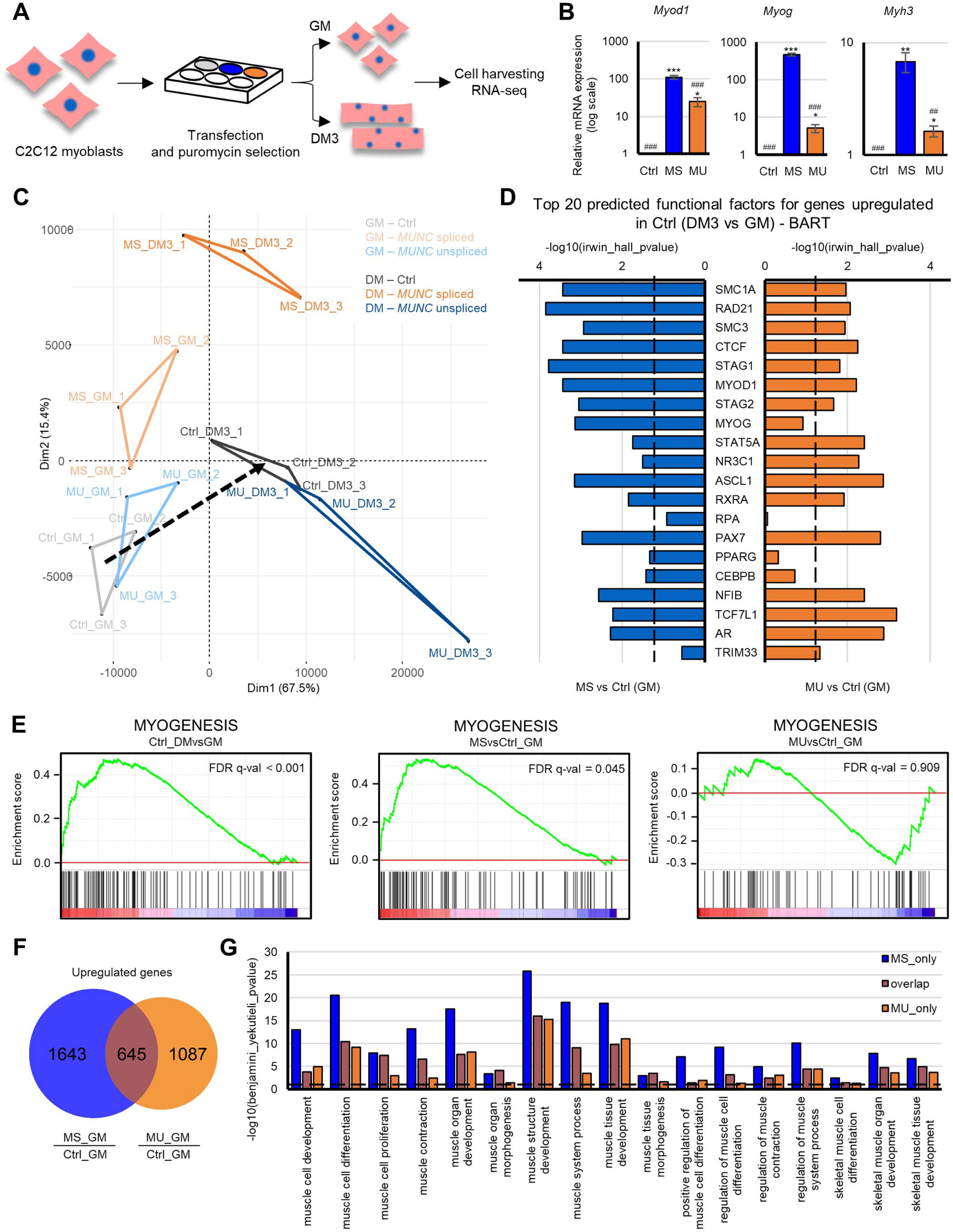
*MUNC* Isoforms Regulate Different Sets of Genes Involved in Promyogenic Pathways. (A) Scheme of experimental design. C2C12 murine myoblasts were transfected with linearized vector with or without *MUNC* overexpression. Grey: control cells (Ctrl); blue: cells overexpressing spliced *MUNC* (MS); orange: cells overexpressing unspliced *MUNC* (MU). Cells were grown in proliferating (GM) or differentiating (DM3) conditions. (B) RT-qPCR analysis of Ctrl, MS, and MU cells grown under proliferating conditions. Levels of mRNAs were normalized to *Gapdh* and are shown relative to Ctrl. Values represent three biological replicates presented as mean ± SEM. Statistical significance was calculated using Student’s t test. *, **, and *** indicate adjusted p < 0.05, 0.01, and 0.001, respectively, in comparison to Ctrl. ## and ### indicate adjusted p < 0.01 and 0.001, respectively, in comparison to MS. (C) Principal component analysis of RNA-seq data. Triangles connect biological replicates from each cell line from specific conditions. Black dashed line indicates differentiation direction. (D) The top 20 transcription factors regulating the upregulated genes in differentiation of Ctrl cells as determined by BART compared with factors for genes upregulated in proliferating MS vs. Ctrl cells (blue) and factors for genes upregulated in proliferating MU vs. Ctrl (orange). Black dashed line represents Irvin-Hall p value of 0.05. (E) Enrichment plots of the gene set involved in myogenesis in differentiating versus proliferating Ctrl cells (Ctrl_DMvsGM), proliferating MS versus Ctrl cells (MSvsCtrl_GM), and proliferating MU versus Ctrl cells (MUvsCtrl_GM). (F) Venn diagram of the overlap between differentially upregulated genes in proliferating MS cells (MS_GM/Ctrl_GM) and differentially upregulated genes in proliferating MU cells (MU_GM/Ctrl_GM). (G) The Benjamini-Yekutieli p values for 17 muscle-related Gene Ontology terms enriched in upregulated genes in proliferating MS or MU cells and the overlap. Black dashed line represents Benjamini-Yekutieli p value of 0.05.

RNA-seq analysis reveals that although the two isoforms are both promyogenic, they induce different sets of genes. Hierarchical clustering of the RNA-seq data confirmed that cells in proliferating conditions were distinct from cells grown in differentiating conditions (Figure S1A). The principal components most responsible for distinguishing proliferating and differentiating expression profiles in control cells are exacerbated upon *MUNC* overexpression, but, intriguingly, the spliced and unspliced isoforms contributed to changes in different axes (Figure 1C). In proliferating conditions, overexpression of either spliced or unspliced *MUNC* pushed cells toward an expression profile more similar to that of differentiated control cells than to control cells grown under proliferating conditions, and cells that overexpressed the spliced isoform had transcriptomes more similar to that of differentiated control cells than did cells that expressed the unspliced isoform. In differentiation medium, when endogenous *MUNC* is expressed, overexpression of the spliced isoform induced more differentiation related genes in the PCA2 direction, whereas the unspliced isoform had a lesser effect in the PCA1 direction (Figure 1C). *MUNC* overexpression altered expression of many genes, although fewer genes responded to *MUNC* overexpression than when cells were simply switched to differentiating conditions (Figure S1B). Thus, *MUNC* overexpression promotes a pro-differentiation gene expression profile, and this phenotypic shift can even be observed in C2C12 in proliferating (non-differentiating) conditions.

To identify the transcription factors that regulate the genes altered by *MUNC* overexpression, we employed Binding Analysis for Regulation of Transcription (BART), which identifies transcription factors enriched in a set of promoters relative to the factors’ genome-wide binding site distribution (Wang et al., 2018). We first identified the 20 transcription factors most highly activated during normal differentiation (Figure 1D). Nineteen of these regulators were also activated by overexpression of at least one *MUNC* isoform in proliferating cells, and fifteen were activated by both (Figure 1D). The top 10 predicted transcription factors are known to contribute to myogenesis (Figure S1C and S1D). The cohesin complex is involved in regulation of genes upregulated by the *MUNC* spliced isoform (Figure S1C), supporting the hypothesis that *MUNC* recruits this complex (Tsai *et al*., 2018).

The *MUNC* isoforms produced distinct changes in gene expression when overexpressed in proliferating cells. Gene Set Enrichment Analysis indicated that the genes upregulated by the spliced isoform in proliferating cells are involved in myogenesis, similar to the enrichments observed between differentiating and proliferating control cells (Figure 1E). Interestingly, overexpression of the unspliced *MUNC* isoform in proliferating cells did not result in significant enrichment for genes associated with myogenesis (Figure 1E), suggesting that either the spliced is simply better than the unspliced isoform at inducing promyogenic genes or that the two isoforms regulate distinct sets of genes. In support of the latter hypothesis, 645 genes were upregulated by both isoforms of *MUNC* (e.g., *Myod1*, *Myog* and *Myh3*; Figure 1B), whereas there were 2730 genes upregulated by one isoform and not the other (Figure 1F and S1E). Gene Ontology (GO) analysis revealed that both non-overlapping and overlapping upregulated gene sets are involved in muscle-related pathways (Figure 1G) and that the genes upregulated by the expression of the spliced isoform were more significantly enriched for muscle-related pathways than those upregulated by expression of the unspliced isoform. The same patterns were observed for genes downregulated in proliferating cells and for genes up- and down-regulated by *MUNC* overexpression in differentiating conditions (Figure S1E). In summary, both isoforms of the *MUNC* lncRNA induce expression of promyogenic genes, likely through activation of a common set of transcription factors. The genes regulated by the two isoforms are distinct, and the spliced isoform has a more pronounced effect on promyogenic and differentiation gene expression profiles than does the unspliced isoform.

### SHAPE-MAP of MUNC Reveals Distinct Secondary Structures

To determine the structural features of *MUNC* lncRNAs and to find the structural modules important for its promyogenic activity, we analyzed the structures of the spliced and unspliced isoforms using SHAPE-MaP chemical probing (Smola *et al*., 2015a; Smola *et al*., 2015b; Weeks, 2021). The RNAs were probed using the SHAPE reagent 5-nitroisatoic anhydride (5NIA) in C2C12 cells (in-cell) and after gentle extraction from C2C12 cells (cell-free). SHAPE measures local nucleotide flexibility, and thus unpaired nucleotides are preferentially acylated at their 2′-hydroxyl groups. SHAPE-modified nucleotides are identified as mutations and short deletions in cDNAs created during relaxed fidelity MaP reverse transcription. The resulting *MUNC* mutation profiles enabled us to model the secondary structures of the spliced and unspliced *MUNC* isoforms.

The SHAPE analysis of the spliced isoform identified multiple intrinsically well-determined regions likely to form well-defined local domains (Figure 2A and 2B). The SHAPE-supported model differs substantially from previously reported *MUNC* lncRNA structures predicted using bioinformatic algorithms (Cichewicz *et al*., 2018). SHAPE reactivities from two independent experiments performed 3 years apart showed good agreement (Pearson’s R = 0.94; Figure S2A) and yielded similar pairing probabilities (Figure S2B). In-cell and cell-free SHAPE data for the *MUNC* spliced isoform are highly correlated (Pearson’s R = 0.92; Figure S2C), suggesting that the in-cell structure is similar to that of the cell-free RNA. We also identified nucleotides with significant in-cell protection from or enhancement of modification relative to the cell-free structure (Figure 2B and S2D), highlighting regions of potential protein interactions or other changes in cells.

**Figure 2.**
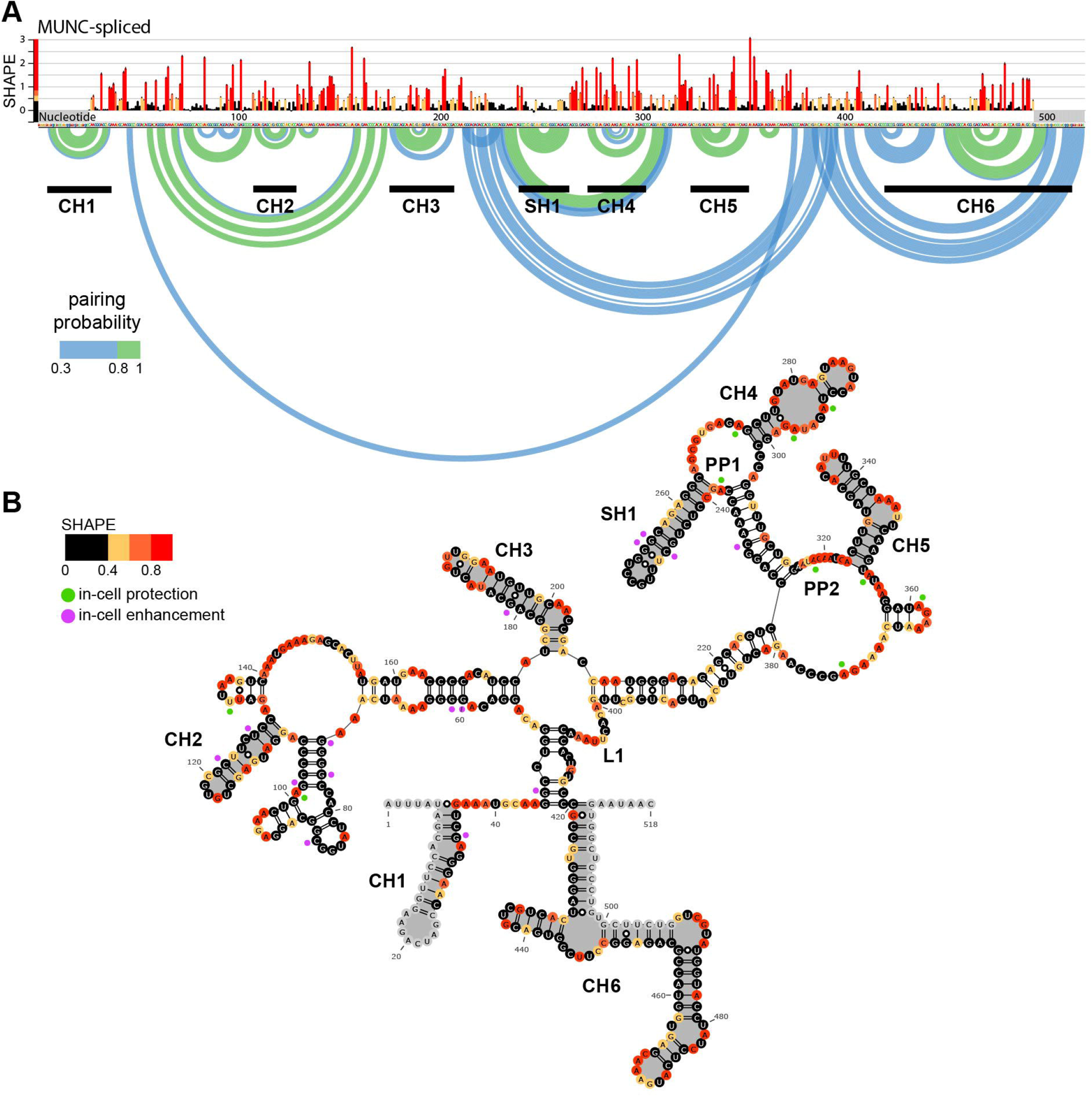
SHAPE-MaP of *MUNC* Spliced Isoform Reveals a Structured Architecture. (A) Top: Nucleotide-resolution SHAPE reactivity profile for the cell-free *MUNC* spliced isoform from two biological replicates. Mean reactivities (± SE) are colored by relative value. Bottom: The pairing probabilities of nucleotide pairs represented by arcs linking the involved nucleotides. Color indicates pairing probability. Domains common to spliced and unspliced *MUNC* (CH1-CH6) and domain specific to the spliced isoform (SH1) are indicated. (B) Minimum free energy secondary structure model of spliced *MUNC* isoform color coded by SHAPE reactivities. In-cell protections are in green and in-cell enhancements are in purple. Structural domains are highlighted in grey.

Comparisons of SHAPE reactivity profiles (Figure S3A) and base-pairing probabilities (Figure S3B) for the spliced isoform (518 nt; exon 1 and exon 2) and the unspliced isoform (1083 nt; exon 1, intron, and exon 2) indicate that the isoforms share six structurally homologous domains (referred to as <u>C</u>ommon <u>H</u>airpin (CH) domains; Figure 2B and S3C). CH1 and CH6 domains are at the 5’ and 3’ ends of the transcripts, respectively, whereas CH2 is close to 3’ end of exon 1. We also discovered regions of well-defined structure unique to the spliced isoform (Figure 2B): one hairpin (SH1), two protein protected sites (PP1 and PP2), and one well-defined loop (L1).

### Distinct Structural Domains of the Spliced Isoform of MUNC Regulate Myogenesis

We next tested how disruption of structural domains in the spliced isoform of *MUNC* identified using SHAPE-based modeling affected the expression of promyogenic factors. A series of variants lacking specific structural domains or containing defined mutations were overexpressed in C2C12 cells (Figure 3A, S4A-C). The wild-type isoform induced production of *Myod1*, *Myog*, and *Myh3* mRNAs in proliferating cells. Deletion or mutation of all tested sites decreased *Myod1* induction, and ΔCH1 and ΔCH4 deletions completely inhibited *Myod1* induction (Figure 3B). *Myog* induction did not require all domains: In mutants with deletions of CH3, CH4, SH1, PP1, and L1, *Myog* induction was observed (Figure 3C). The motifs dispensable for *Myog* induction were dispensable for *Myh3* induction as well with the exception of CH4 (Figure 3D). Notably, in the ΔCH1 mutant induction of none of the three promyogenic factors was detected (Figure 3A-3D), suggesting that CH1 plays a critical role in activity of the spliced isoform of *MUNC*. Overexpression of the wild-type spliced isoform in differentiating cells also induced the three promyogenic factors, and ΔCH1 and ΔCH4 impaired in induction of all three promyogenic factors in differentiating cells (Figure S4D-F).

**Figure 3.**
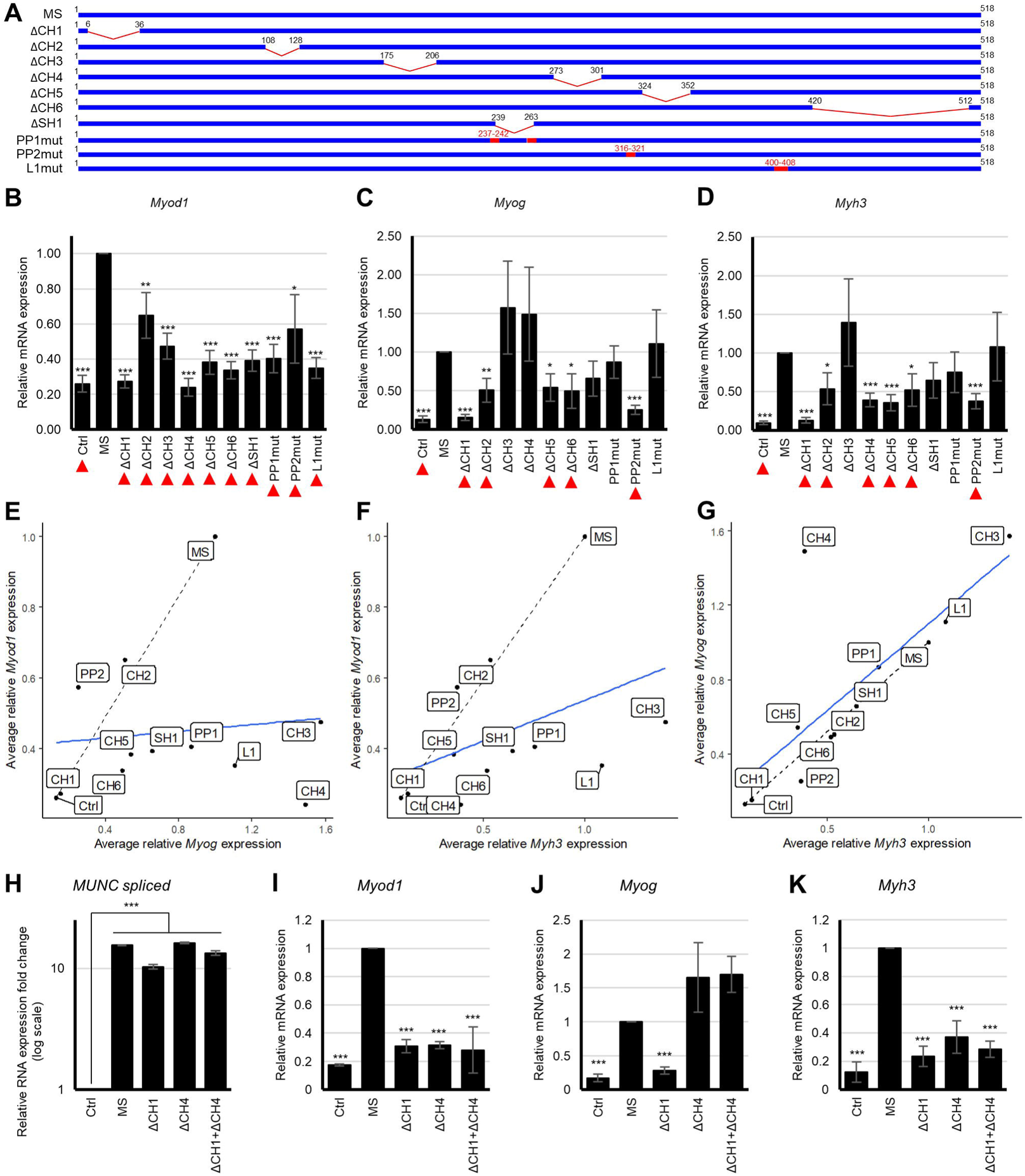
Distinct Structural Domains of the Spliced Isoform of *MUNC* are Required for Induction of Promyogenic Factors. A) Deletion and mutation constructs used in the study. (B-D) RT-qPCR analyses of B) *Myod1*, C) *Myog*, and D) *Myh3* mRNAs in extracts of proliferating control cells (Ctrl), cells that express the wild-type spliced isoform of *MUNC* (MS), and cells that express domain mutants. Data were normalized to *Gapdh* and are shown relative to MS. Values represent at least two independent transfectants with at least three biological replicates for each of them and are presented as means ± SEM. Statistical significance was calculated using the Student’s t test. ***, **, and * indicate p < 0.001, <0.01, and < 0.05, respectively, in comparison to MS. Red arrows indicate significantly different results. (E-G) Plots of average induction of E) *Myod1* vs. *Myog*, F) *Myod1* vs. *Myh3*, and G) *Myog* vs. *Myh3* in Ctrl cells and cells that express MS and domain mutants. Black dashed line indicates theoretical correlation line if the expression of one gene is dependent on another. Blue line shows experimental correlation between analyzed mRNAs upon expression of indicated constructs. (H) RT-qPCR analyses of *MUNC* constructs in proliferating cells. Levels of lncRNAs were normalized to *Gapdh* and are shown relative to Ctrl. Values represent three biological replicates and are presented as means ± SEM. Statistical significance was calculated using Student’s t test. *** indicates p < 0.001 in comparison to Ctrl. (I-K) RT-qPCR analyses of levels of I) *Myod1*, J) *Myog*, and K) *Myh3* mRNAs in cells expressing the indicated MUNC constructs. Levels were normalized to *Gapdh* and are shown relative to MS. Values represent three biological replicates and are presented as means ± SEM. Statistical significance was calculated using Student’s t test. *** indicates p < 0.001 in comparison to MS.

The mutations that disrupted CH4, CH3, and L1 impaired *Myod1* induction but promoted expression of *Myog*, suggesting that the lncRNA activates *Myog* independently of *Myod1* in proliferating cells (Figure 3E). Different *MUNC* domains also influenced *Myh3* and *Myod1* expression (Figure 3F). Mutations in domains of spliced *MUNC* affected *Myh3* and *Myog* expression similarly, except ΔCH4 that had a deleterious effect on *Myod1* and *Myh3* expression but not *Myog* expression (Figure 3G). These correlations suggest that *MUNC* induces *Myog* independent of *Myod1* while *Myh3* induction is mostly dependent on *Myog* with some additional dependence on *Myod1*.

In some cases, functional domains can cooperate when supplied either *in cis* or *in trans* (Uroda et al 2019), so we tested whether ΔCH1 and ΔCH4 variants complement each other without being physically connected. Co-overexpression of ΔCH1 and ΔCH4 *in trans* did not rescue defects in *Myod1* or *Myh3* induction in proliferating (Figure 3H-K) or differentiating cells (Figure S4G-J). We conclude that the local proximity of CH1 and CH4 elements on the same RNA molecule is important for induction of expression of *Myod1* and of *Myh3*.

### Overexpression of MUNC Leads to Phenotypical Changes in C2C12 Cells

Overexpression of the spliced isoform of *MUNC* induced production of MYOD1 protein in proliferating conditions and differentiation conditions (Figure 4A-B, S5A-B). Phenotypical changes and an increase in the frequency of MHC-positive cells were also observed upon overexpression of the spliced isoform of the lncRNA (Figure 4C-D). Induction of MYOD1 protein was not observed in proliferating cells when the *MUNC* construct lacked CH1, CH4, or CH5 domains (Figure 4B). In differentiation conditions, ΔCH1 and ΔCH4 did not induce MYOD1 protein expression, although other constructs did (Figure S4A-B). Although *Myog* and *Myh3* mRNA are induced by expression of the wild-type spliced isoform in proliferating conditions, MYOG and MHC protein expression required differentiation.

**Figure 4.**
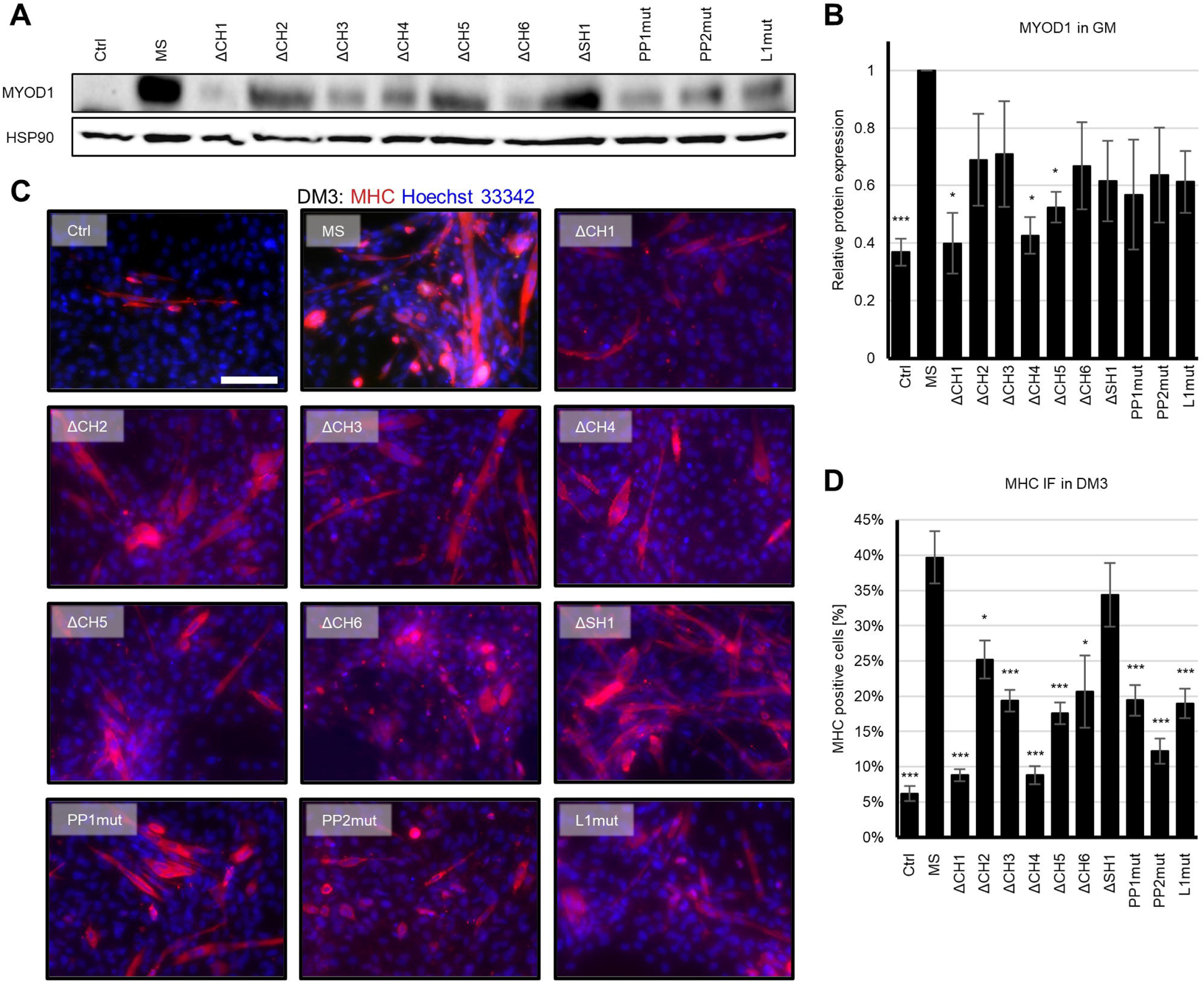
*MUNC* Structural Domains are Necessary for Myotube Formation. (A) Representative Western blot for MYOD1 in proliferating control cells (Ctrl) and cells overexpressing the wild-type spliced *MUNC* (MS) or mutants overexpressing cells. HSP90 served as a loading control. (B) Quantification of MYOD1 protein in proliferating cells that express indicated *MUNC* constructs normalized to HSP90 and shown relative to MS. Values represent two independent transfectants with two biological replicates for each of them and are presented as means ± SEM. Statistical significance was calculated using Student’s t test. *** and * indicate p < 0.001 and < 0.05, respectively, in comparison to MS. (C) Representative immunofluorescence images of fixed cells that express indicated *MUNC* constructs 3 days after differentiation (DM3). Cells were immunostained with antibodies against MHC. Hoechst 33342 was used to visualize nuclei. Scale bar, 200 µm. (D) Quantification of the percentage of MHC-positive cells in cultures of cells that express the indicated *MUNC* constructs. Values represent two independent transfectants with at least 1000 nuclei counted and are presented as means ± SEM. Statistical significance was calculated using Student’s t test. *** and * indicate p < 0.001 and < 0.05, respectively, in comparison to MS.

Given that protein levels are affected by *MUNC* spliced overexpression in domain-specific manner, we tested C2C12 differentiation efficiency as measured by percentage of MHC-positive cells upon overexpression of the various *MUNC* constructs. Expression of the wild-type spliced *MUNC* isoform increased the percentage of MHC-positive cells at 3 days of differentiation (Figure 4C-D). The effect of *MUNC* lncRNA on differentiation was especially dependent on CH1 and CH4 domains (Figure 4C-D).

### MUNC Transcripts Lacking CH1 and CH4 are Structurally Similar to Wild-type

When performing structure-function studies by making targeted deletions in RNAs, it is important to assess whether local sequence perturbations lead to extensive structural consequences. To ensure that the loss of phenotype observed upon deletion of CH1 or CH4 domains and lack of complementation *in trans* are not due to misfolding, we performed cell-free SHAPE-MaP on wild-type *MUNC* spliced, ΔCH1 and ΔCH4 constructs. The SHAPE-informed secondary structure models and reactivity data of ΔCH1 and ΔCH4 are highly similar to each other and to the wild-type spliced isoform (Figure 5A-C). The reactivity data for these two constructs were highly correlated with the data for the wild-type spliced isoform (Pearson’s R values 0.93 and 0.94, respectively, Figure 5D and 5E). These results suggest that global lncRNA structure is essentially the same in all cases. The structures of all other structural domains and conserved helices are preserved in both ΔCH1 and ΔCH4 mutants. These data, first, support the overall accuracy of our structural models, as specific well-defined motifs can be deleted without affecting the global structure. Thus, we conclude that CH1 and CH4 domains are critical for the observed promyogenic phenotypes.

**Figure 5.**
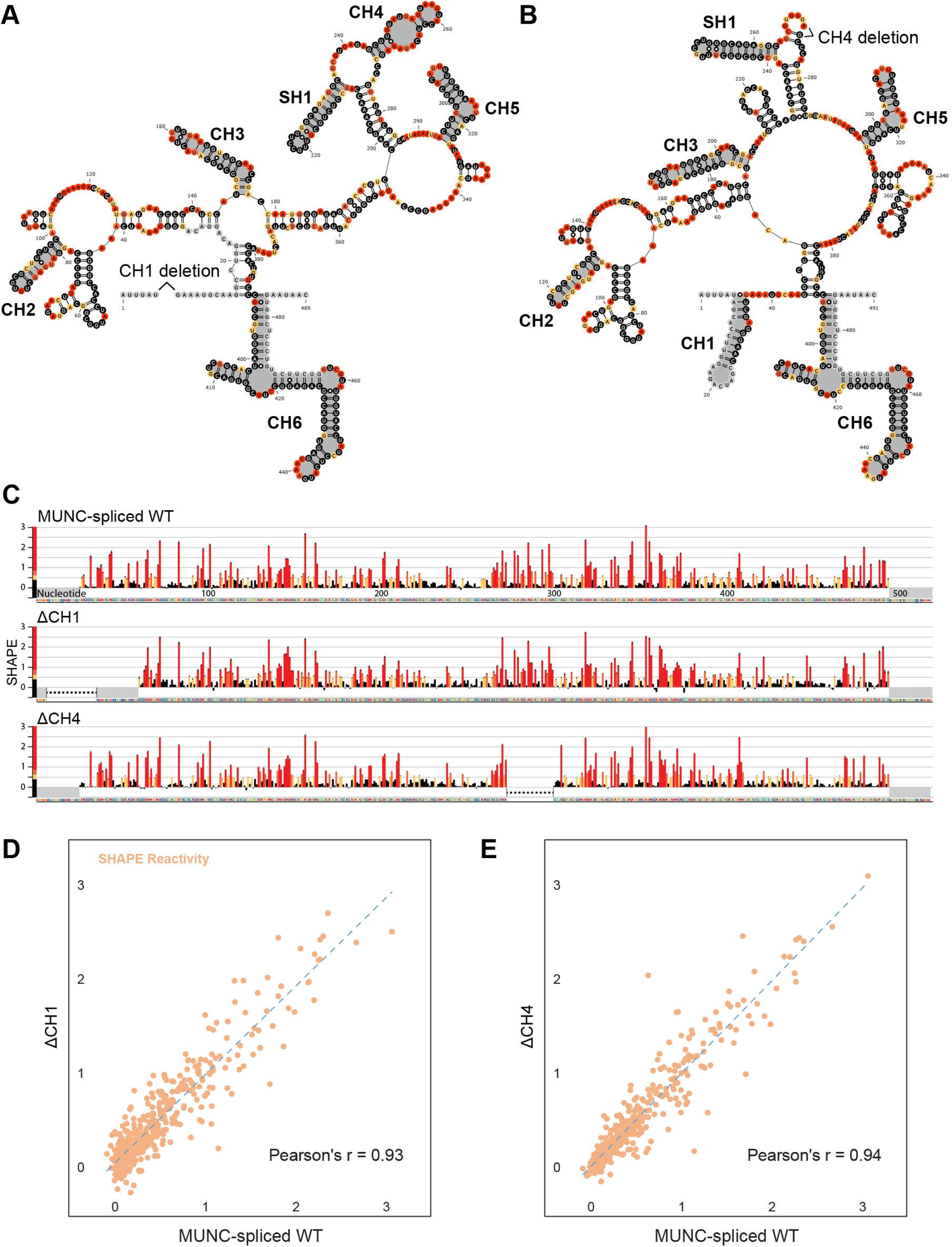
Deletion of CH1 or CH4 Domains Does Not Disrupt *MUNC* lncRNA Structure. (A) Minimum free energy secondary structure model of ΔCH1 color coded for SHAPE reactivities. Structural domains are highlighted in grey. (B) Minimum free energy secondary structure model of ΔCH4 color coded for SHAPE reactivities. Structural domains are highlighted in grey. (C) Nucleotide-resolution SHAPE reactivity profile for the wild-type spliced isoform, ΔCH1, and ΔCH4 from cell-free SHAPE-MaP. Mean reactivities (± SE) are colored by relative value. Grey boxes indicate primers used for PCR. Black dashed lines represent deletions. (D) Correlation plot of SHAPE reactivities for wild-type (x axis) and ΔCH1 (y axis) spliced isoforms. (E) Correlation plot of SHAPE reactivities for wild-type (x axis) and ΔCH4 (y axis) spliced isoforms.

### MUNC Binds to Specific Genomic Sites and Regulates Expression of Adjoining Genes in a Domain-Specific Manner

We next investigated whether *MUNC* physically associates with the genes it regulates (Figure 6A). Using the Chromatin Isolation by RNA Purification (ChIRP) assay we identified *MUNC* binding sites at 410 genomic loci. More than 90% of identified targets overlap with previously published results (Tsai *et al*., 2018) (Figure S6A), and the locations of binding sites relative to transcription start sites of nearby genes were very similar between the two data sets (Figure S6B-C). To understand whether *MUNC* lncRNA binding to chromatin affects gene expression regulation, we compared our ChIRP-seq targets to RNA-seq profiles of wild-type cells in proliferating and differentiating conditions to those of cells that overexpress *MUNC* and to expression microarray data from cells where *MUNC* RNA has been depleted (Mueller *et al*., 2015). Hierarchical clustering demonstrated that the expression profiles of *MUNC*-targeted genes more closely resembles the expression in proliferating conditions than differentiating conditions when *MUNC* is depleted (Figure S6D). This confirmed that *MUNC* is required for expression changes during normal myogenesis. In support of the hypothesis that *MUNC* binding to a promoter region directly regulates expression of the downstream gene, overexpression of *MUNC* in proliferating cells results in expression profiles of *MUNC*-bound genes similar to the profile in differentiating conditions (Figure S6E). These results support *MUNC*’s role as a promyogenic factor and that the *MUNC* binding sites are functional and important for skeletal muscle differentiation. We also noted that approximately equal numbers of genes adjoining *MUNC* binding sites are up- or downregulated after *MUNC* overexpression and that the genes are distributed over all chromosomes and not clustered around the *MUNC* locus (Figure 7A). Thus, *MUNC* can act *in trans*, and can activate or repress genes that are proximate to sites where it is bound.

**Figure 6.**
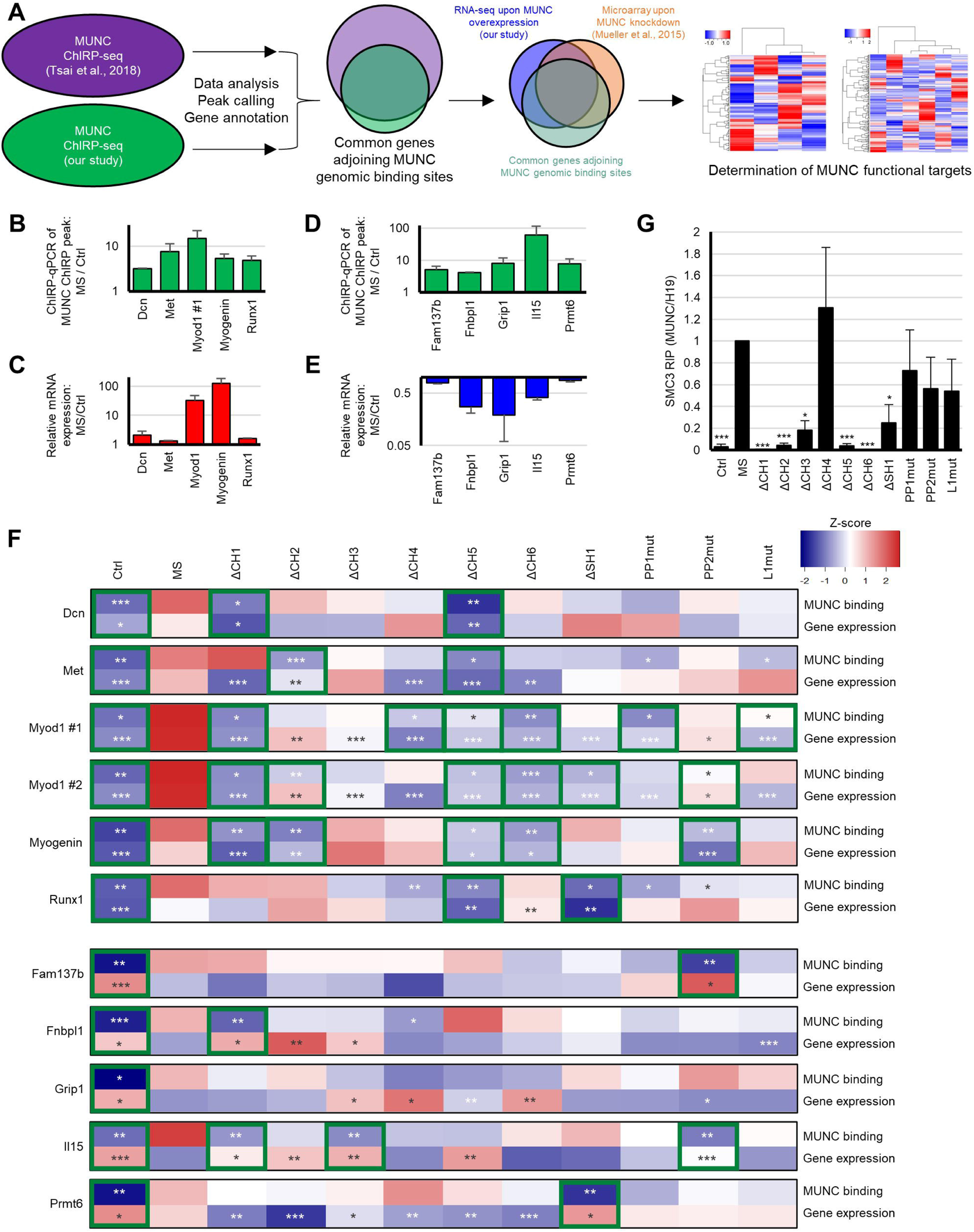
Distinct *MUNC* Spliced Domains Are Required for Binding to Specific Genomic Sites, Regulating Adjoining Genes Expression, and Interacting with Protein Partners. (A) Scheme of ChIRP-seq analyses followed by overlap with the RNA-seq data after *MUNC* overexpression and microarray data after *MUNC* knockdown to identify functional *MUNC* targets. (B, D) ChIRP-qPCR of *MUNC* genomic binding sites (ChIRP peaks). Values from three biological replicates are presented as mean ± SEM from *MUNC* spliced overexpressing cells over control cells. Statistical significance shown in (F). (C, E) RT-qPCR analysis of expression of genes adjoining to ChIRP peaks. Values from three biological replicates are presented as mean ± SEM from *MUNC* spliced overexpressing cells over control cells. Statistical significance shown in (F). (F) Heatmap of ChIRP-qPCR (“*MUNC* binding”) and RT-qPCR (“Gene expression”) Z-scores for 11 analyzed ChIRP peaks and corresponding 10 adjoining genes (top: genes upregulated upon *MUNC* spliced overexpression, bottom: genes downregulated upon *MUNC* spliced overexpression) for proliferating control cells (Ctrl), *MUNC* spliced wild-type overexpressing cells (MS) and *MUNC* spliced mutants overexpressing cells (ΔCH1-L1mut). Mean values from three biological replicates are presented as Z score. Color scale presented in right upper corner. ***, **, * indicates p-value < 0.001, <0.01, and < 0.05, respectively, in comparison to MS. Green boxes presents domains required both for MUNC binding and gene expression regulation. (G) SMC3 RIP-qPCR results for proliferating control cells (Ctrl), *MUNC* spliced wild-type overexpressing cells (MS) and *MUNC* spliced mutants overexpressing cells (ΔCH1-L1mut). Levels of *MUNC* lncRNA were normalized to *H19* lncRNA and shown relative to MS. Values represent three biological replicates and are presented as mean ± SEM. Statistical significance was calculated using Student’s t test. ***, * indicates p-value < 0.001 and < 0.05, respectively, in comparison to MS.

**Figure 7.**
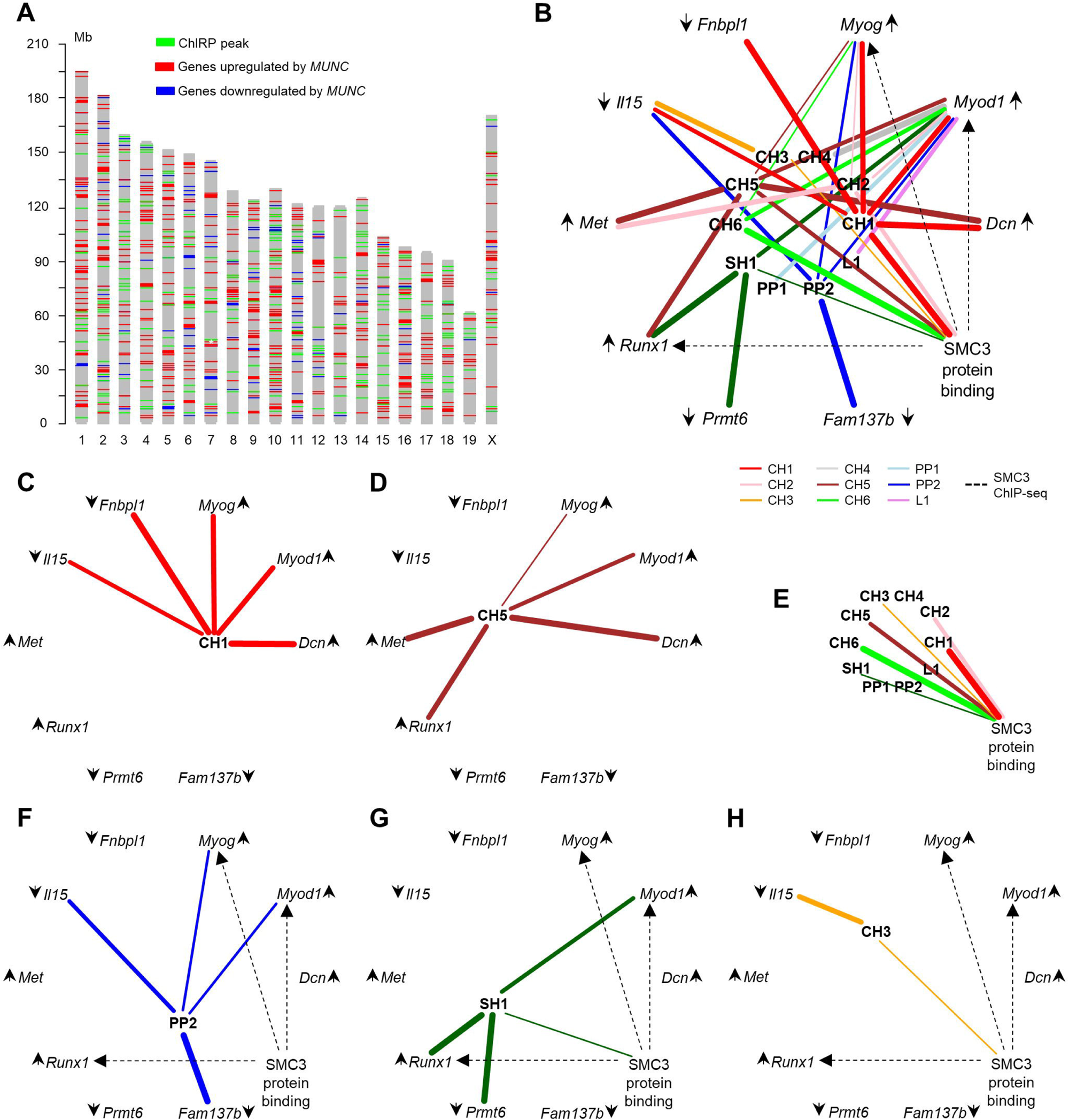
*MUNC* Regulates Genes *in Cis* and *in Trans* through Multiple Mechanisms. (A) Map of chromosomal locations of *MUNC* ChIRP peaks (green), genes upregulated by *MUNC* spliced overexpression (red), and genes downregulated by *MUNC* overexpression (blue). White star marks *Myod1* locus. (B) Graph summarizing all interactions identified in this study. Lines represent significant alterations relative to overexpression of wild-type spliced *MUNC* phenotype (DNA binding and adjoining gene expression regulation or SMC3 protein binding) upon deletion/mutation of particular domain. Line width represents effect size. Arrows next gene names show the effect of *MUNC* (upregulation or downregulation). The dashed lines represent SMC3 binding sites in promoter regions of *Myod1*, *Myog*, and *Runx1*. (C-G) Graphs as in panel B summarizing C) interactions of CH1, D) interactions of CH5, E) domains required for optimal *MUNC*-SMC3 binding, F) interactions of PP2, and G) interactions of CH3.

To establish which of the *MUNC* targets are important for myogenesis, we separately evaluated genes activated and genes repressed by *MUNC*. The *MUNC-*activated genes were defined as *MUNC*-bound genes that are induced during normal differentiation, repressed by depletion of *MUNC*, and induced by *MUNC* overexpression. The *MUNC*-repressed genes were defined as *MUNC*-bound genes that are repressed during normal differentiation, induced by *MUNC* depletion, and repressed by *MUNC* overexpression. There were 37 *MUNC*-activated genes and 22 direct *MUNC*-repressed genes that are regulated in a concordant manner during myogenesis (Figure S6F). ChIRP-qPCR for 10 of them showed that *MUNC* binding to the sites adjoining these genes was significantly increased after *MUNC* overexpression, and RT-qPCR confirmed significant up- or downregulation of these genes (Figure 6B-6E).

We next evaluated the effects of *MUNC* mutants on binding to the promoter regions and on gene expression (Figure 6F and Figure S7). In particular, the CH1 domain is critically important for *MUNC* binding and activation of *Myod1, Myog*, and *Dcn* and repression of *Fnbpl1* and *Il15*. The CH5 domain is important for *MUNC*-mediated regulation of *Myod1, Myog, Dcn, Met*, and *Runx1*. Although all domains were required for *Myod1* induction by *MUNC* (Figure 2B), specific domains are important for binding to the two *MUNC*-binding sites in the *Myod1* promoter region. CH4, PP1, and L1 are important for binding to one site, and CH2, SH1, and PP2 are important for binding to the other; CH1, CH5, and CH6 are required at both sites. In another example, PP2 is required for *MUNC* to bind to and downregulate the *Fam137b* gene, whereas CH1 is required for downregulation of *Fnbpl1*. Thus, different *MUNC* structural domains regulate different sets of target genes. In addition, binding to and regulation of some genes require the function of multiple domains (Figure 7B). This argues against a common mechanism of *MUNC* function on all target genes.

*MUNC* is known to interact with the cohesin complex to regulate gene expression (Tsai *et al*., 2018). We thus expected that certain domains of *MUNC* would be important for binding to SMC3, a key component of the cohesin complex. RNA-immunoprecipitation with an anti-SMC3 antibody and RIP-qPCR of *MUNC* demonstrated that deletions of CH1, CH2, CH3, CH5, CH6, or SH1, but not CH4, PP1, or PP2, significantly decreased *MUNC* binding to SMC3 (Figure 6G). Together these data show that *MUNC* regulates gene expression through distinct domains and that multiple mechanisms may be important for DNA/protein binding and gene regulation (Figure 7B).

## Discussion

Analyses of protein structure to define domains responsible for specific functions has proven essential for understanding complex functions. Analogous structure-directed functional studies for lncRNAs are rare. Here, we performed a structure-function analysis of the two isoforms of the *MUNC* lncRNA, which function during differentiation of skeletal muscle myoblasts (Mueller *et al*., 2015). Critically, the secondary structures determined based on modeling directed by SHAPE-MaP data differed substantially from those predicted using RNA folding algorithms (Cichewicz *et al*., 2018; Tsai *et al*., 2018), emphasizing the need for experimental-directed structural studies and modeling of lncRNAs (Weeks, 2021).

We performed a structure-guided analysis of the importance of various domains that are common to the two *MUNC* isoforms and that are unique to the spliced isoform. The mechanism of action of *MUNC* lncRNA is more complex than expected for an eRNA, as different RNA domains of *MUNC* are important for binding to and regulating different genes. Importantly, we showed that the mutations made to either delete entire domains or to specifically disrupt structure in a particular domain were surgical in nature and did not alter the overall structure of the lncRNA.

Both the spliced isoform and the unspliced isoform promote myogenesis and share multiple well-defined structural motifs but also contain isoform-specific structural domains. Although the two isoforms differ by the presence of a single intron, they clearly have non-identical effects. The two isoforms do activate a common subset of genes, as well as a larger non-overlapping set of genes, and both sets are enriched in muscle related pathways. The spliced transcirpt is also a much stronger promyogenic factor than unspliced isoform. Multiple factors might influence the two isoforms divergent behaviors. First, the spliced isoform is more stable structurally: it has a larger fraction of well-determined (low entropy) structures than the unspliced transcript. Second, different sites on the two isoforms may be bound by varying cellular factors: the relative changes in reactivity profiles between in-cell and cell-free conditions, a measure of interactions with cellular factors, are much higher for the spliced than unspliced isoform. Because most structural domains in spliced isoform also exist within unspliced one, we hypothesize that the intron in unspliced transcript binds to additional factors that allow it to target different genes. Lastly, the included intron in unspliced isoform may contribute more to *Myod1* transcription *in cis* when functioning as an eRNA, while the more stably folded regions which assemble after *MUNC* splicing could regulate other targets *in trans*.

To evaluate the mechanisms by which the spliced isoform of *MUNC* regulates gene expression, we focused on *Myod1*, *Myog*, and *Myh3*, which encode promyogenic transcription factors. We previously reported that *MUNC* overexpression leads to increases in levels of these mRNAs (Cichewicz *et al*., 2018; Mueller *et al*., 2015). By careful comparison of effects of different *MUNC* mutants, we showed that *MUNC* directly regulates expression of all three mRNAs, although its effect on *Myh3* appears to be MYOG- and MYOD1-dependent in proliferating murine myoblasts. This observation confirms *MUNC* function is not limited to the secondary effects of MYOD1 expression resulting from *MUNC* enhancer RNA activity.

To interrogate whether different regions of *MUNC* act independently or whether they must be *in cis*, we focused on CH1 and CH4. These two domains are the most important of the domains tested for *Myod1* induction, and deletion of either did not affect the structure of the other as shown by SHAPE-MaP analyses. The mutants with deletions of CH1 and of CH4 did not complement each other *in trans* to rescue *Myod1* induction. Therefore, CH1 and CH4 motifs must be exist on the same molecule to stimulate *Myod1* expression *in trans*.

Our structural and functional studies revealed that *MUNC* has multiple effector domains and does not regulate expression all its target genes by a common mechanism. Because *MUNC* could be recruiting multiple transcription complexes, we focused on genes that are directly regulated by *MUNC* binding near their promoter. We used ChIRP to determine the sites in the genome where *MUNC* binds and looked for cases where *MUNC* also regulates adjoining gene expression to find potentially direct targets of *MUNC*. Distinct combinations of *MUNC* domains are required for binding to and induction of *Dcn* (CH1 and CH5), *Met* (CH2 and CH5), and *Runx1* (CH5 and SH1), genes that are known to stimulate muscle differentiation (Kishioka et al., 2008; Li et al., 2020b; Umansky et al., 2015; Wang et al., 2005). Similarly, different domains were critical for downregulation of *Il15* (CH1, CH3, and PP2)*, Prmt6* (SH1), and *Grip1* (CH4), genes that are known to have negative impact on skeletal muscles (Choi et al., 2019; Quinn et al., 1995; Wu et al., 2005). Thus, the multi-modal mechanisms of regulation of gene expression by *MUNC* involve different domains of *MUNC*, and likely different protein partners. *MUNC* recruits varying cooperative machinery depending on the genomic context, and this behavior is likely exemplary of other lncRNAs.

It was previously proposed that MUNC regulates expression of *Myog* by recruiting the cohesin complex to its promoter (Tsai *et al*., 2018). We confirmed that *MUNC* binds to SMC3 and then showed that CH1, CH2, CH3, CH5, CH6, and SH1 domains of *MUNC* are specifically required for this interaction. SMC3 binds in the promoter regions of *Myod1, Myog*, and *Runx1* (Tsai *et al*., 2018); however, *MUNC*-SMC3 binding is not sufficient for all *MUNC*-mediated gene expression regulation. For example, mutation of the PP2 domain did not change *MUNC*-SMC3 binding but decreased induction of *Myod1* and *Myog* by *MUNC* (Figure 7F). Conversely, although *MUNC*-SMC3 binding is diminished upon deletion of SH1 or CH3, *MUNC* can still induce *Myog* (Figure 7G), and *Myod1, Myog* and *Runx1* (Figure 7H), respectively. Thus, in some cases *MUNC*-SMC3 binding may not be sufficient or required for regulating gene expression by *MUNC*.

In conclusion, this study established the power of integrating experimentally-driven secondary structure modeling with structure-function analyses to identify functional domains and mechanisms of action of lncRNAs. Indeed, all domains chosen for validation based on our models proved to be functional. Our study clarifies the role of RNA structure in *MUNC* function. *MUNC* was initially described as a *cis*-acting eRNA for *Myod1* and was soon thereafter shown to have *trans*-acting functions beyond those expected for a classical eRNA (Cichewicz *et al*., 2018; Mueller *et al*., 2015; Tsai *et al*., 2018). We show that this underlying functional complexity of *MUNC* is integrated with its RNA structural complexity. Our results confirm *MUNC* regulation of *Myod1* and complex regulation of other genomic targets *in trans*. In addition, our study shows that binding of SMC3, and by association cohesin, is not sufficient for *MUNC* function, and may not be essential for regulating all of the target genes such as *Myod1*, *Myog* and *Runx1*.

The compact and highly organized structure of *MUNC* lncRNA differentiates it from longer lncRNAs like *Xist* and *HOTAIR,* that present long unstructured repetitive sequence regions landing pads for protein multimerization (Smola *et al*., 2016; Wang et al., 2017). *MUNC* instead appears to have scaffolding functions more similar to those of *ANRIL* (Zhang et al., 2020). *MUNC* may act as an eRNA *in cis* to recruit a regulatory protein complex (like cohesin) as it does for *Myod1* or *in trans* as demonstrated for *Myog* and *Runx1*. *MUNC* binding can also inhibit some target gene transcription as for *Il15*, *Prmt6*, and *Grip1* by binding to their promoter region and diminishing activity of a transcriptional regulatory factor; in these cases, *MUNC* may act as a decoy or may directly occlude a binding site. Lastly, since *MUNC* is encoded 5-kb upstream of the transcription start site of *Myod1*, the transcription of *MUNC* itself may positively regulate the transcription of *Myod1* and other loci in close 3-dimensional proximity by maintaining active chromatin structure. *MUNC* may also play a role in stabilizing the genome organization and control the spreading of posttranslational modifications to nearby chromatin. Regardless, experimentally derived structure models are essential for rapid characterization of functionally important motifs within RNAs of interest.

## Acknowledgments

We thank Genome Analysis and Technology Core at University of Virginia (Charlottesville, USA) for RNA-seq and ChIRP-seq sequencing. This work was supported by the grants from the NIH (R01 AR067712 to AD, R35 GM122532 to KMW, and R35-GM128635 to MJG), an American Cancer Society Postdoctoral Fellowship (ACS 130845-RSG-17-114-01-RMC to CAW), a Predoctoral Fellowship from the American Heart Association (18PRE33990261 to RKP), Wagner Fellowships from the University of Virginia (to RKP and MAC), and the F99/K00 NCI Predoctoral to Postdoctoral Fellow Transition Award (F99CA253732 to RKP).

## Author Contributions

RKP conceived of the project and, together with AD, designed the experiments. CAW and KMW designed, analyzed, and interpreted all SHAPE experiments. PSI performed all SHAPE-MaP sequencing. RKP and PP performed all other bioinformatic analyses with help and guidance from MJG. MAC prepared *MUNC* ChIRP-seq libraries. RKP cloned all *MUNC* mutant vectors with help of KNJ. SS performed SMC3 RIP-qPCR experiment, which was designed and analyzed by RKP. RKP performed RNA-seq, SHAPE-MaP, differentiation assays, RT-qPCR, Western blots, MHC immunofluorescence, and ChIRP-qPCR. RKP, CAW, KNJ, and AD wrote the manuscript with input from all authors. All authors reviewed and edited the manuscript and approved the final draft.

## STAR Methods text

### RNA-seq

RNA samples were isolated from proliferating or differentiating control cells or cells that overexpress a *MUNC* construct by TRIzol extraction using Direct-zol RNA MiniPrep Plus Kit including DNase treatment. RNA-seq was performed by Hudson Alpha on poly(A)-enriched RNA using the Illumina HiSeq 2500 instrument. RNA-seq data was aligned to the mouse assembly GRCm38/mm10 using STAR v2.5 (Dobin et al., 2013) and quantified by HTSeq (Anders et al., 2015). DESeq2 R package (Love et al., 2014) was then applied to identify differentially expressed genes with a adjusted p of <0.05. Bioinformatic prediction for functional factors (including transcription factors and chromatin regulators) that bind at *cis*-regulatory regions was performed using BART 2.0 (Wang *et al*., 2018). Gene set enrichment analysis was performed as previously described (Subramanian et al., 2005). Gene Ontology was performed using GeneTrail2 (Stöckel et al., 2016). All RNA-seq library data files are available under GEO accession number GSE174203 as a part of the SuperSeries GSE174218.

### SHAPE-MaP

#### Cell-free SHAPE

Control or *MUNC* construct overexpressing C2C12 cells were grown to 70% confluency in two 15-cm dishes. Both plates were washed once in PBS before scraping and lysis in 2.5 mL of proteinase K buffer (40 mM Tris, pH 8, 200 mM NaCl, 1.5% sodium dodecyl sulfate, and 0.5 mg/mL proteinase K). Proteins were digested for 45 min at 23 °C with intermittent mixing. Nucleic acids were extracted twice with 1 volume of phenol:chloroform:isoamyl alcohol (25:24:1) that was pre-equilibrated with 1.1× RNA folding buffer (110 mM HEPES, pH 8, 110 mM NaCl, 5.5 mM MgCl_2_). Excess phenol was removed through two subsequent extractions with 1 volume chloroform. The final aqueous layer was buffer exchanged into 1.1 × RNA folding buffer using PD-10 desalting columns (GE Healthcare Life Sciences). The resulting RNA solution was incubated at 37 °C for 20 min before being split into two equal volumes. The SHAPE reagent, 250 mM 5NIA (AstaTech) in DMSO was added to one half, and DMSO was added to the other. Samples were incubated at 37 °C for 10 min. RNA was precipitated with 1/10 volume of 2 M NH_4_OAc and 1 volume of isopropanol. After one wash with 75% ethanol, the resulting pellet was dried and resuspended in 88 μL of water and 10 μL of 10× TURBO DNase buffer and 4 units of TURBO DNase (Thermo Fisher) were added. The mixture was incubated at 37 °C for 1 h. RNA was purified (GeneJET RNA Cleanup and Concentration Micro Kit, Fisher) and eluted into 20 μL of nuclease-free water.

#### In-cell SHAPE

Control C2C12 cells or cells expressing *MUNC* constructs were grown to 70% confluency in four wells of a 6-well plate. After washing with PBS, 900 μL of standard growth medium was added. Next, 100 μL of 250 mM 5NIA was added to two wells and 100 μL of DMSO (control) were added to the other two wells. Plates were incubated at 37 °C for 10 min. Media was aspirated, cells were washed once with PBS, and total RNA was extracted using TRIzol (Thermo Fisher). RNA pellets were dried and resuspended in 88 μL nuclease-free water, treated with TURBO DNase, and purified with GeneJET RNA Cleanup and Concentration Micro Kit as described for the cell-free experiment.

#### MaP Reverse Transcription

1 μg of each RNA sample was subjected to MaP reverse transcription, which requires Superscript II and addition of betaine and Mn^2+^ to the RT buffer (Siegfried et al., 2014; Smola *et al*., 2015b), using a *MUNC*-specific reverse primer (Table S1). The cDNA generated was buffer exchanged over Illustra microspin G-50 columns (GE Healthcare). For second-strand cDNA synthesis, output DNA (corresponding to 167 ng of total RNA) was used as a template for 25 μL PCR reactions (Q5 Hot-start polymerase, NEB) with primers made to amplify spliced (1-518 bp) and unspliced (1-792 and 584-1083 bp) *MUNC* isoforms. Reactions included 1x Q5 reaction buffer, 250 nM each primer, 100 μM dNTPs, 0.02 units/μL Q5 Hot-start polymerase. PCR was conducted as follows: 98 °C for 30 s, then 25 cycles of 98 °C for 10 s, 60 °C (*MUNC* spliced) or 69 °C (*MUNC* unspliced) for 30 s, and 72 °C for 35 s, followed by 72 °C for 2 min. Step 1 PCR products were run on a 2% gel and purified using Zymoclean Gel DNA Recovery Kit and eluted in 10 μL of nuclease-free water. Purified PCR products were measured using a Qubit dsDNA HS Assay Kit, and 1 ng was used for tagmentation using Nextera XT DNA Library Preparation Kit (Illumina). After neutralization with NT buffer, multiplex indices were added using the Nextera XT DNA Library Preparation Kit (Illumina). PCR was performed as follows: 72 °C for 3 min, 95 °C for 30 s, then 12 cycles of 95 °C for 10 s, 55 °C for 30 s, 72 °C 30 s, and, finally, 72 °C for 5 min. Step 2 PCR products were purified using a 0.8x ratio of Agencourt AMPure XP beads (Beckman Coulter) and eluted in 20 μL of nuclease-free water.

#### Sequencing of MaP libraries

Size distributions and purities of fragmented *MUNC* amplicon libraries were verified (2100 Bioanalyzer, Agilent). Libraries (about 120 amol of each) were sequenced on a MiSeq instrument (Illumina) with 2 × 250 or 2 × 300 paired-end sequencing. Libraries derived from total cytoplasmic RNA were sequenced with 2 × 300 paired-end sequencing on a MiSeq instrument, combining reads from multiple runs until desired RNA sequencing depth was achieved. All SHAPE-MaP libraries data files are available under GEO accession number GSE174217 as a part of the SuperSeries GSE174218.

#### Mutation counting and SHAPE profile generation with ShapeMapper 2 software

FASTQ files from sequencing runs were directly input into the ShapeMapper 2 software (Busan and Weeks, 2018) for read alignment and mutation counting. To ensure that mutation rates were not affected by reduced fidelity at reverse transcription initiation sites, target FASTA files input to ShapeMapper 2 had primer-overlapping sequences and the first 5 nucleotides transcribed were set to lowercase, which eliminates these positions from analysis. ShapeMapper 2 was run with the --min-depth 4000 flag and all other values set to defaults. In each experiment, the 5NIA-treated samples were designated as the “modified” samples and DMSO-treated samples as “unmodified” samples.

#### Modeling *MUNC* structure using SuperFold

The SuperFold analysis software (Smola *et al*., 2015b) was used with experimental SHAPE data to inform RNA structure modeling by RNAStructure (Reuter and Mathews, 2010). Default parameters were used to generate base-pairing probabilities for all nucleotides (with a max pairing distance of 600 nt) and minimum free energy structure models.

#### Identification of in-cell changes in *MUNC* spliced SHAPE reactivity

SHAPE reactivities of in-cell and cell-free treated RNAs were normalized to each other using a median difference minimization strategy. First, the log relative reactivities for each dataset were calculated as follows:

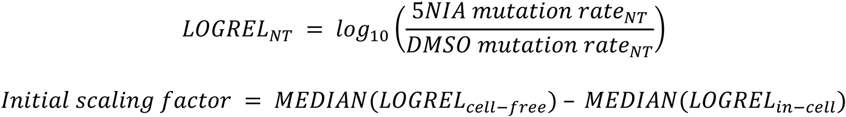

The LOGREL_in-cell_ values were adjusted up by the initial scaling factor, and differences were calculated for each nucleotide:

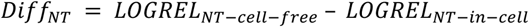

The final scaling factor (added to in-cell LOGREL values) was calculated as the value that minimizes the median for all nucleotides of | Diff_NT_|. New Diff_NT_ values were computed with the final scaling factor, and Z-scores were computed for each nucleotide:

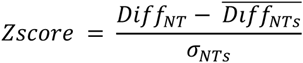

Only nucleotides with Z-scores > 1.645 standard deviations from the mean (90% confidence intervals) in both replicates were considered significant shifts in SHAPE reactivity.

#### Graphical display of SHAPE data

Secondary structure projection images were generated using the (VARNA) visualization applet for RNA (Darty et al., 2009).

### Structural analyses of *MUNC* mutants

Cells overexpressing the wild-type spliced *MUNC* isoform and ΔCH1 and ΔCH4 mutants were used for cell-free SHAPE as described above. MaP reverse transcription was performed using a *MUNC*-specific reverse primer. For second-strand cDNA synthesis we used the same reverse primer for all three *MUNC* constructs but a different forward primer was used for ΔCH1 that overlapped the deletion junction. All the further steps were performed as described above.

#### Cell lines generation

The pLPCX plasmid carrying the sequence of spliced *MUNC* (Mueller *et al*., 2015) was used as a template to obtain mutants via PCR followed by In-Fusion cloning. The constructs were linearized and transfected into the C2C12 cells using Lipofectamine 3000 (Life Technologies). After 24 h, pools of stably transfected cells were selected with 2 μg/ml puromycin. The procedure was repeated at least twice for each construct to ensure that the observed phenotype was not an effect of random selection of a less or more differentiation-potent population. For complementation experiments ΔCH4 was cloned into pLHCX vector. At 24 h after transfection, pools of stably transfected cells were selected with 300 μg/ml hygromycin. Oligonucleotide sequences are listed in Table S1.

#### Cell culture and differentiation

For proliferating conditions, C2C12 cells were cultured in DMEM-high glucose medium (GE Healthcare Life Sciences co.) with 20% fetal bovine serum (Gibco). For differentiation, the serum was 2% horse serum (GE Healthcare Life Sciences co.).

#### RNA isolation and RT-PCR

RNA was isolated by TRIzol extraction using Direct-zol RNA MiniPrep Plus Kit including DNase treatment. cDNA synthesis for mRNA expression levels measurement was performed using GoScript RT cDNA synthesis kit (VWR) with random hexamer priming. After cDNA synthesis, qPCR was performed with StepOnePlus™ Real-Time PCR System using PowerUp™ SYBR® Green Master Mix (Thermo Fisher). All primers used in this study are listed in Table S1.

#### Western blotting

Cells were lysed in IPH buffer (50 mM Tris-Cl, 0.5% NP-40%, 50 mM EDTA). Samples were run on a 10% polyacrylamide SDS-PAGE gel and transferred to nitrocellulose membranes. Membranes were blocked for 30 min in 5% milk containing PBST and incubated overnight with primary antibody in 1% milk. After washing, secondary antibody incubation was carried out for 1 h at 1:4000 dilution before washing and incubation with Millipore Immonilon HRP substrate. Antibodies used were as follows: MYOD1 sc-32758 (Santa Cruz Biotechnology), MHC 22287-1-AP (Proteintech), MYOGENIN sc-12732 (Santa Cruz Biotechnology), and HSP90 sc-13119 (Santa Cruz Biotechnology).

#### Immunofluorescence assay

Cells were plated on glass coverslips and collected after 3 days of differentiation. The coverslips were fixed with 4% paraformaldehyde in PBS for 15 min, permeabilized in 0.5% Triton X-100 in PBS, and blocked in 5% goat serum. The coverslips were incubated with primary antibody MHC 22287-1-AP (Proteintech) overnight at 4 °C and then with Alexa Fluor 555-conjugated secondary antibody (Thermo Fisher Scientific) for 1 h. Cells were stained with Hoechst 33342 (1 μg/mL; Invitrogen) for 2 min at room temperature, washed, and then mounted with ProLong Gold (Invitrogen). The primary and secondary antibodies were diluted 1:400 and 1:1000, respectively. Microscopy was performed using the Zeiss Axio Observer Live Cell microscope and ImageJ Software for analysis (Schneider et al., 2012).

#### ChIRP-seq and ChIRP-qPCR

Antisense probes complementary to the genomic *MUNC* sequence (Table S1) were labeled with biotin-16dUTP (Roche, 11093070910) using a terminal transferase reaction (NEB, M0315). Probes were purified with the QIAquick Nucleotide Removal Kit (QIAGEN, Cat: 28304). ChIRP was performed as described (Chu et al., 2012). ChIRP libraries were prepared using DNA SMART ChIP-Seq kit (Takara Bio., 634865) with 1 ng of DNA as starting material. The quality and quantity of final libraries were assessed using an Agilent Technologies 2100 Bioanalyzer. Libraries were sequenced on an Illumina MiSeq platform (Genome Analysis and Technology Core, University of Virginia School of Medicine). All ChIRP-seq libraries data files are available under GEO accession number GSE174195 as a part of the SuperSeries GSE174218.

ChIRP-seq data from our study and from Tsai et al. (Tsai *et al*., 2018) were independently analyzed. First, data was aligned to the mouse assembly GRCm38/mm10 using bowtie2 version 2.3.4.1 (Langmead and Salzberg, 2012). Peak calling was done using MACS2 version 2.1.1.20160309 (Zhang et al., 2008) with a q value cutoff of 0.05. Peaks were assigned to gene-centric genomic regions with GREAT (McLean et al., 2010). To ensure detection of only true positives, we designed an additional set of biotinylated probes based on the structure of the spliced *MUNC* (Table S1). Next, we performed ChIRP followed by qPCR with StepOnePlus™ Real-Time PCR System using PowerUp™ SYBR® Green Master Mix (Thermo Fisher). Oligonucleotide sequences are listed in Table S1.

#### RIP-qPCR

The RNA immunoprecipitation assay (RIP) was carried out as previously described (Klattenhoff et al., 2013) with slight modification. Briefly, 1×10^7^ proliferating cells from each cell line were lysed with lysis buffer, incubated on ice for 20 min, and centrifuged at 2,500xg for 10 min. The nuclear pellet was lysed with RIP lysis buffer (25 mM Tris, pH 7.4, 150 mM KCl, 1 mM DTT, 0.5% NP-40, 1 mM PMSF, 10 mM NaF, 0.25% sodium deoxycholate, phosphatase inhibitor (Roche), protease inhibitor (Thermo Fisher), and 100 U/ml RNase inhibitor) and incubated on ice for 30 min. The cell lysates were sonicated for a total of 30 s (10 s on, 10 s off) with 10% amplitude. The lysates were centrifuged at 10,000xg for 15 min, and 1 mg of cell lysates were incubated with 4 µg of SMC3 antibody (Abcam) at 4 °C overnight. The immuno-complexes were captured with 30 µl Protein G Plus Agarose beads (Thermo Fisher) and incubated at 4 °C for 2 h. The SMC3-bound RNA-protein complexes were washed three times with RIP lysis buffer. Next, 1 ml of TRIzol was added directly to the pellet, and RNA was precipitated with ethanol and glycogen followed by cDNA synthesis and qPCR analysis. Fold enrichment was calculated by taking the ratio of *MUNC* enrichment in SMC3 immunoprecipitated over a negative control long noncoding RNA H19.

#### Other bioinformatic analyses

A graph that summarizes all interactions presented in this study was created using the rTRM package for R (Diez et al., 2014). A map of chromosomal locations was created with the idiogramFISH package for R (Roa F, 2021).

