## Supplementary figures and images for "Distinct *MUNC* lncRNA structural domains regulate transcription of different promyogenic factors"

### Supplementary Figure

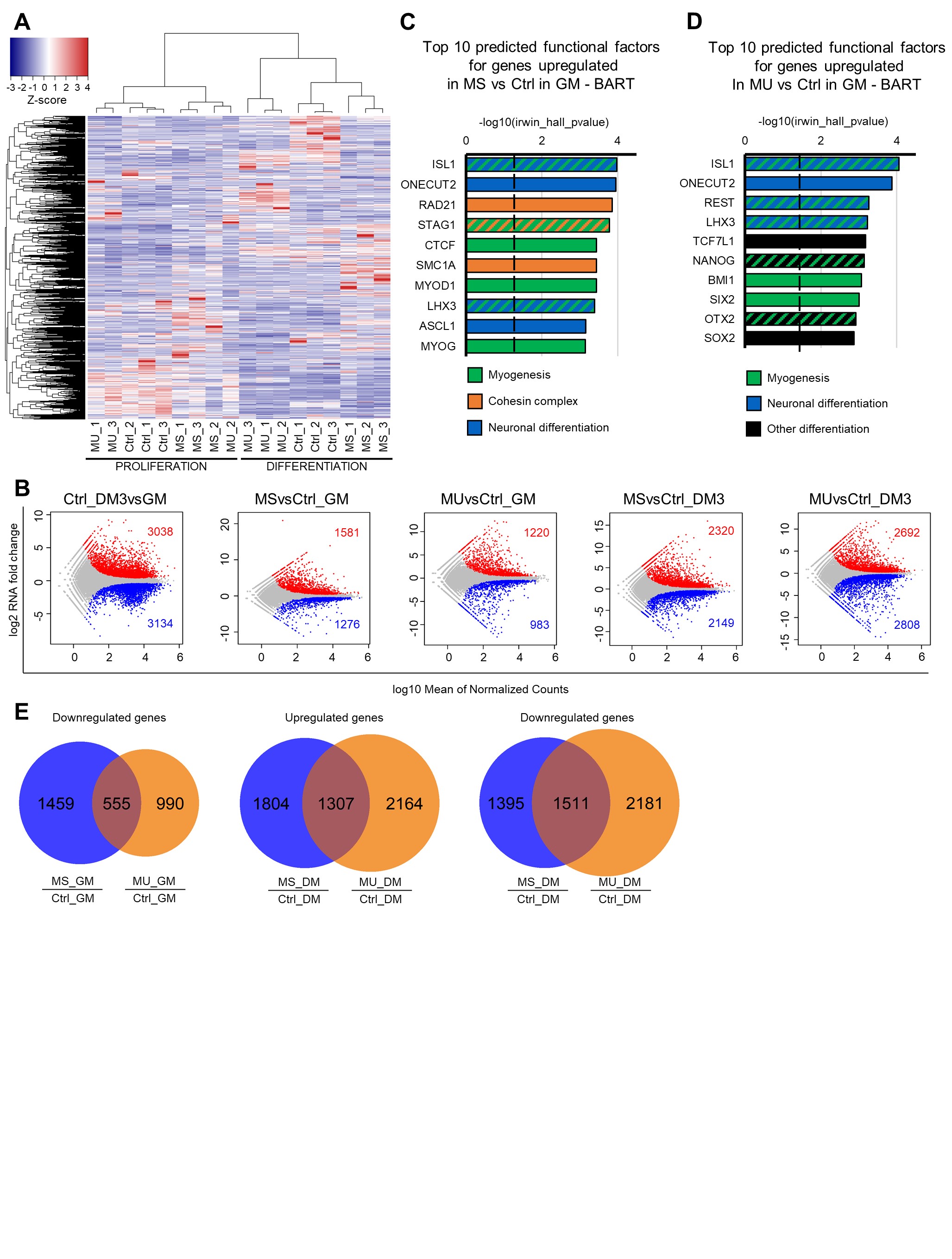

### Supplementary Figure

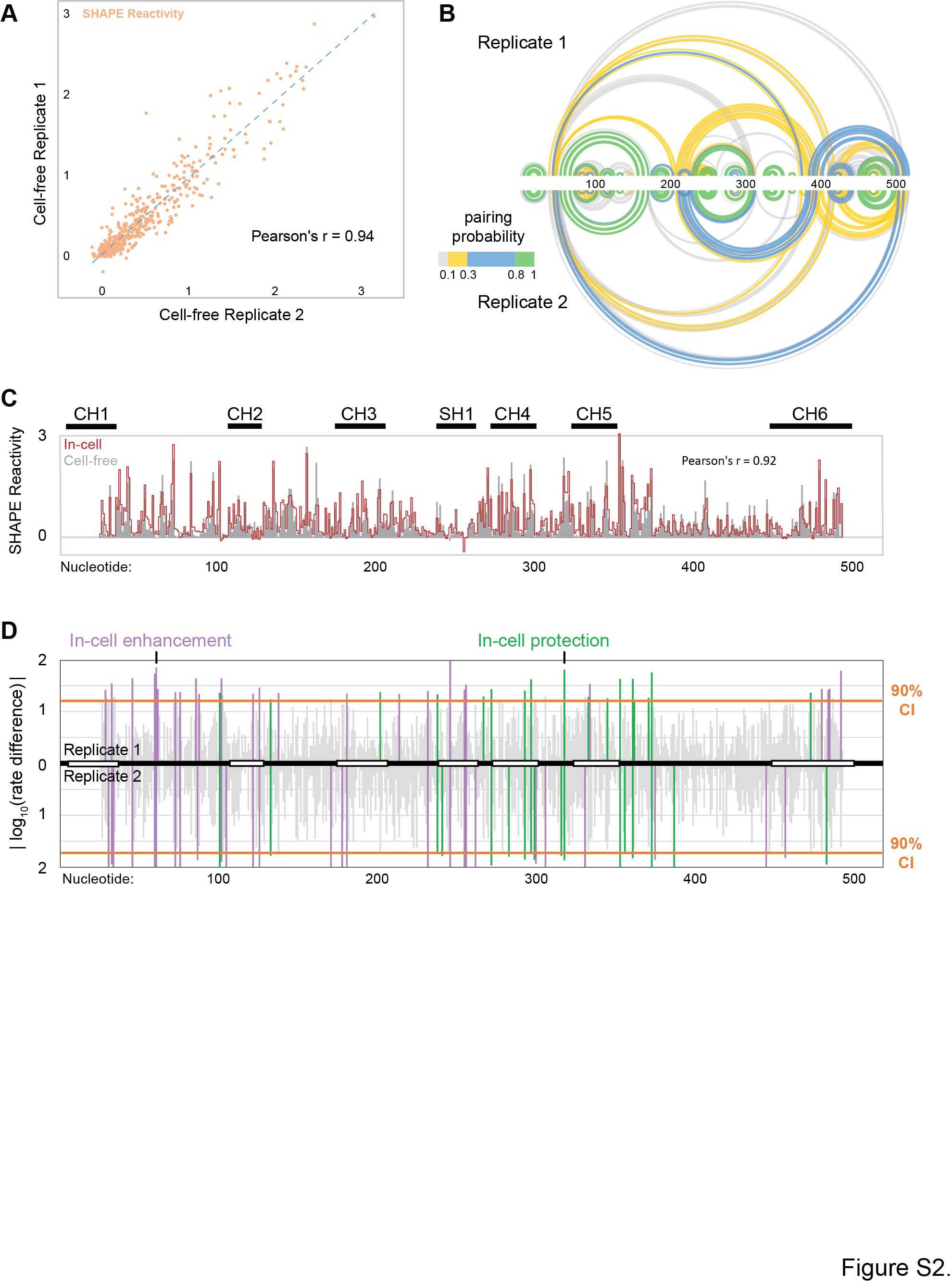

### Supplementary Figure

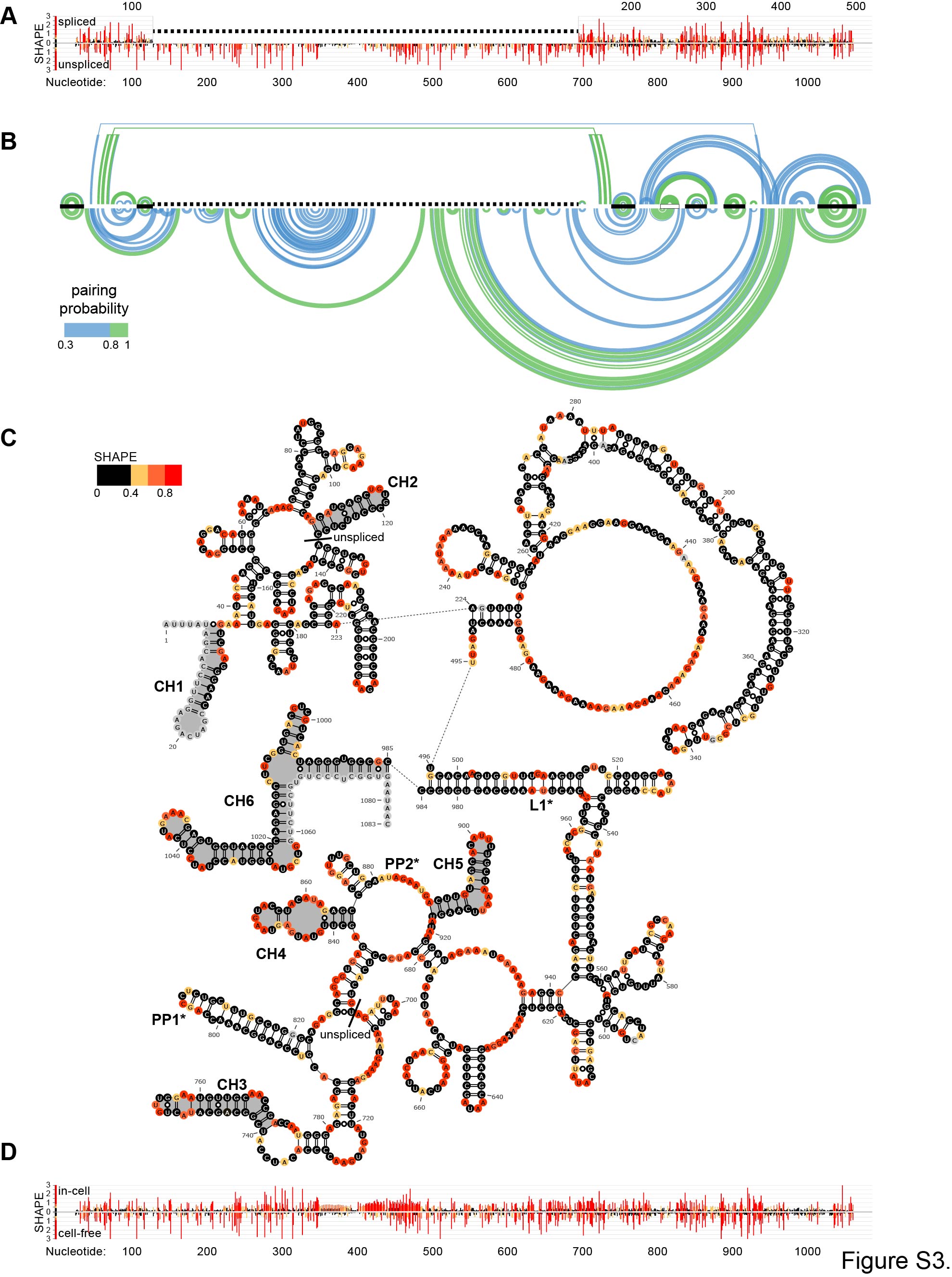

### Supplementary Figure

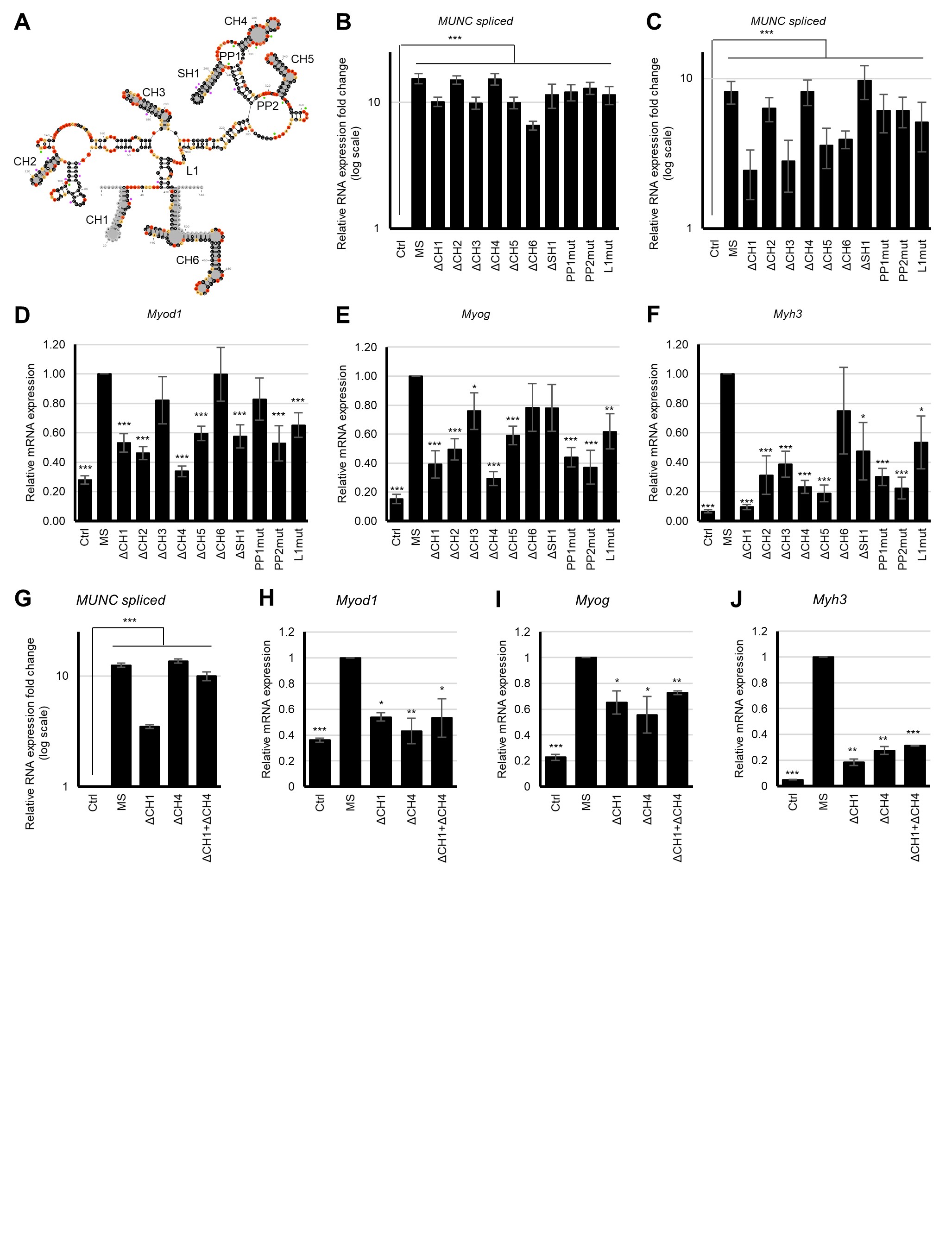

### Supplementary Figure

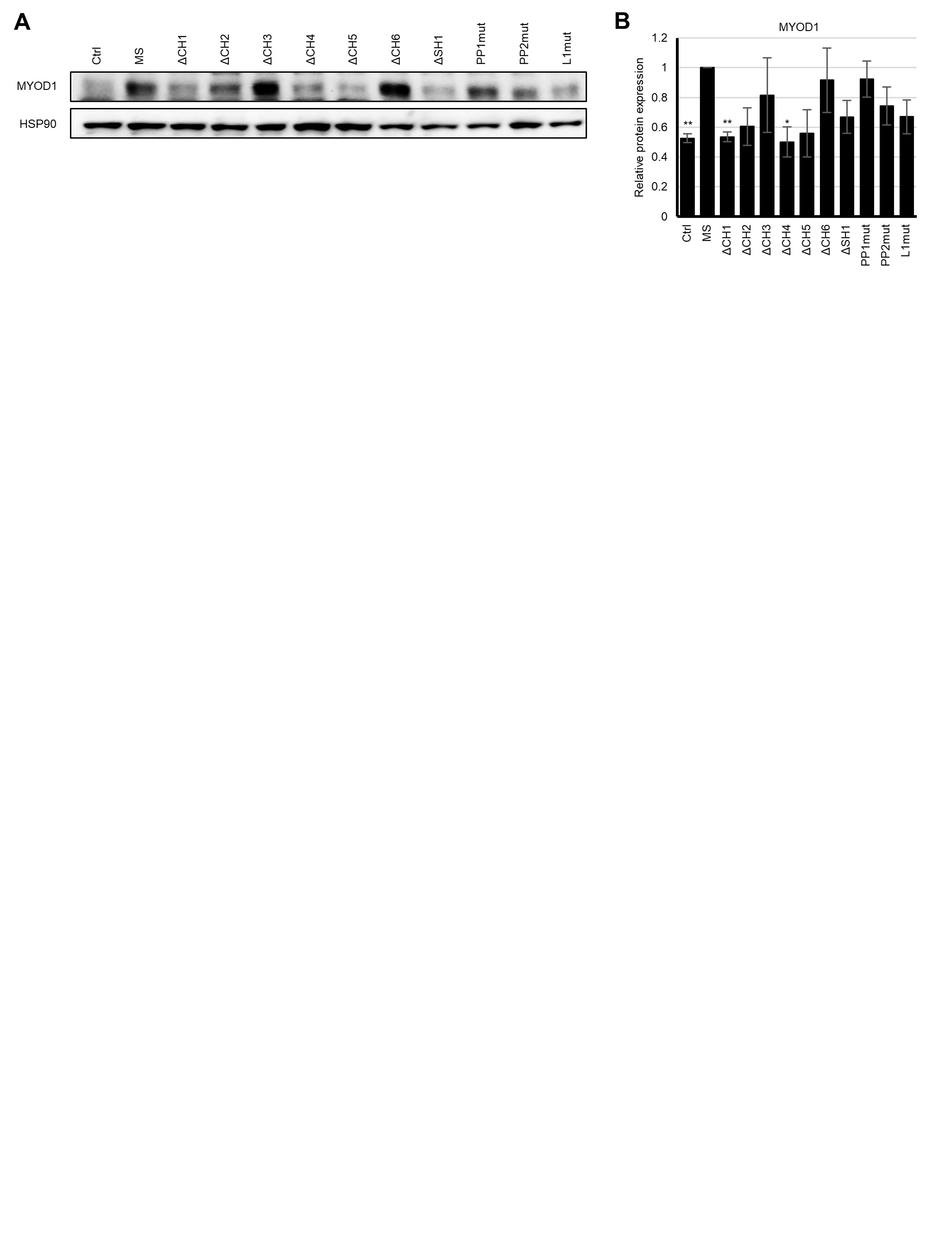

### Supplementary Figure

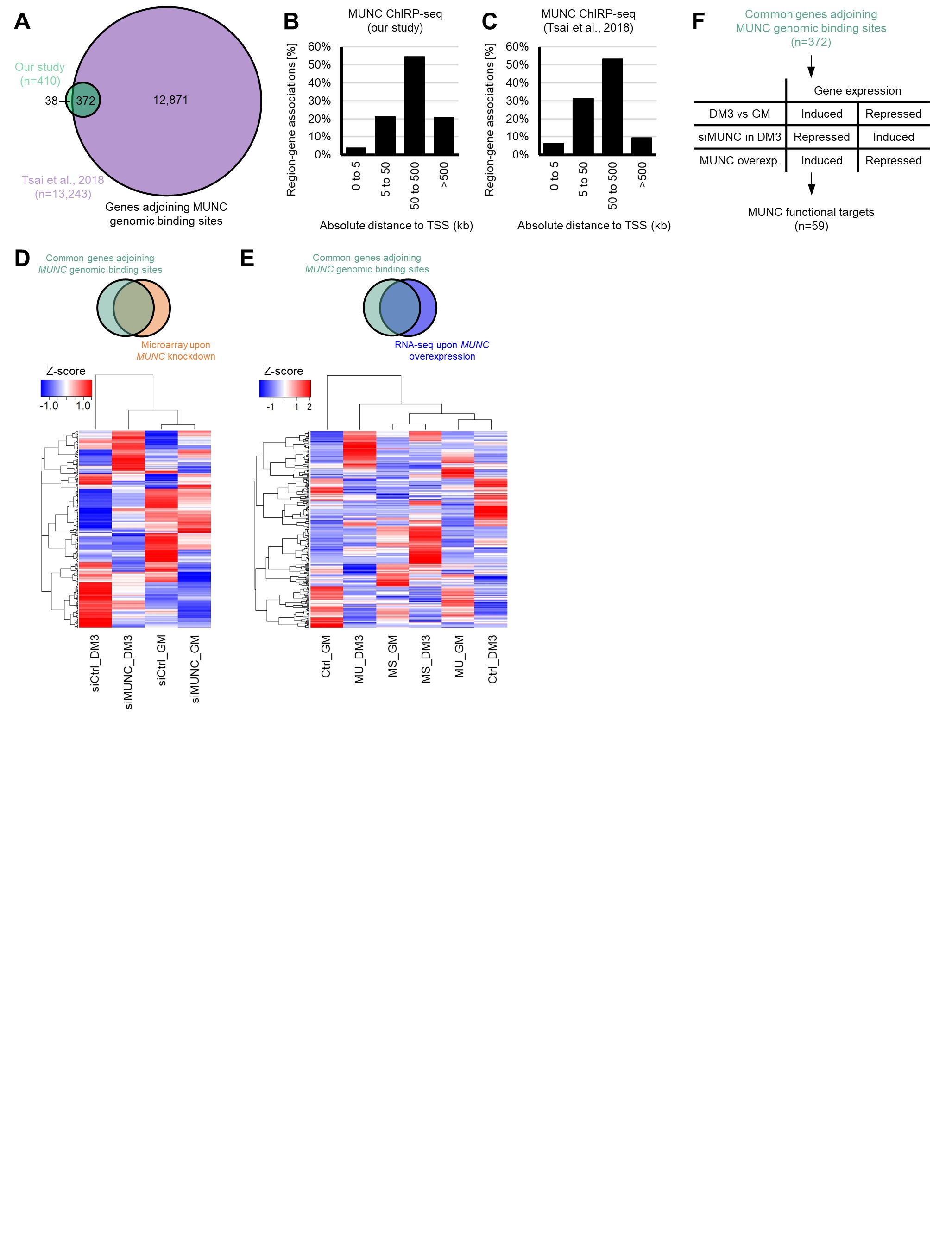

### Supplementary Figure

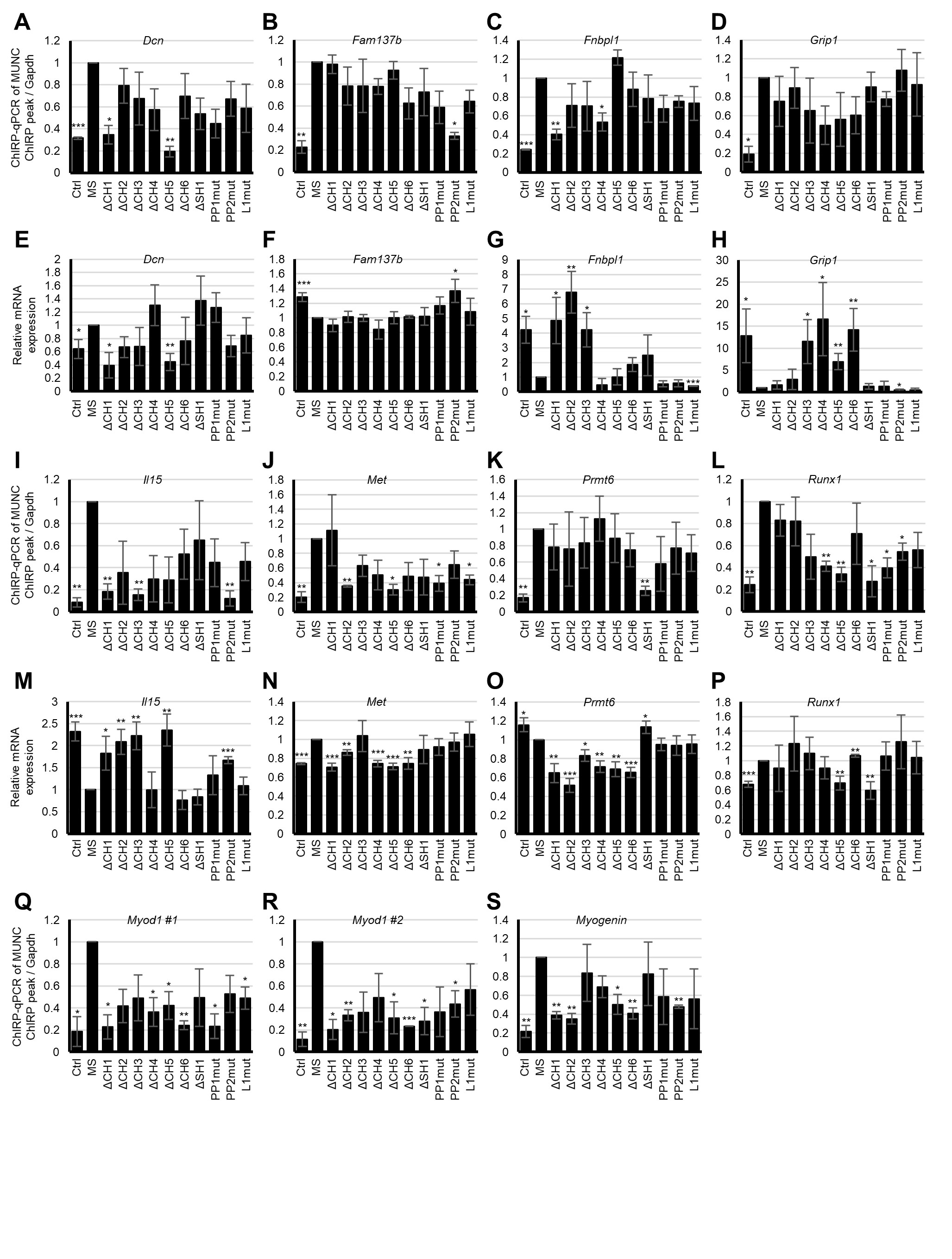
