## Supplementary Figures for "Distinct *MUNC* lncRNA structural domains regulate transcription of different promyogenic factors"

**Supplemental Figure Legend**

*Supplemental Figure 1*. The Spliced Isoform is More Promyogenic than Unspliced *MUNC.*

(A) Heatmap showing clustering of RNA-seq data from all samples and conditions based on expression in proliferating conditions (left) and in differentiating conditions (right) in control cells (Ctrl), cells overexpressing the spliced isoform (MS), and cells overexpressing the unspliced isoform (MU). Numbers indicate the three biological replicates for each condition.

(B) Plots of fold change vs. mean of normalized counts for genes differentially expressed between control cells in differentiating (DM3) and proliferating (GM) conditions, for MS vs. Ctrl in GM, for MU vs. Ctrl in GM, for MS vs. Ctrl in DM3, and for MU vs. Ctrl in DM3. Upregulated genes are in red and downregulated genes are in blue (adjusted $p<0.01$).

(C) BART for genes upregulated in MS vs. Ctrl under proliferating conditions. Black dashed line represents Irvin-Hall p value of 0.05.

(D) BART for genes upregulated in MU vs. Ctrl under proliferating conditions. Black dashed line represents Irvin-Hall p value of 0.05.

(E) Venn diagrams representing overlap between differentially regulated genes in cells that overexpress MS vs. MU.

*Supplemental Figure 2.* Secondary Structure of the *MUNC* Spliced Isoform.

(A) Correlation plot between cell-free SHAPE reactivities from two biological replicates of analysis of the spliced *MUNC* isoform in cell-free conditions.

(B) The pairing probabilities of nucleotide pairs of the spliced *MUNC* isoform in cell-free conditions represented by arcs linking the involved nucleotides. Arc color scale indicates pairing probability.

(C) In-cell and cell-free SHAPE reactivity profiles of spliced *MUNC*. Domains detected in both spliced and unspliced *MUNC* are labeled CH1-CH6. The spliced isoform-specific domain is labeled SH1.

(D) In-cell enhancements and protections based on comparison of in-cell and cell-free SHAPE reactivity profiles of spliced *MUNC* from two biological replicates. White boxes indicate locations of structural domains CH1-CH6 and SH1.

*Supplemental Figure 3*. Comparison of SHAPE-MaP-Derived Structures of Spliced and Unspliced Isoforms Reveals Common Domains.

(A) Nucleotide-resolution SHAPE reactivity profiles for spliced (top) and unspliced (bottom) cell-free RNAs. Mean reactivities (± SE) are colored by relative value. Black dashed line represents intron.

(B) Pairing probabilities of nucleotide pairs indicated by arcs linking the involved nucleotides for the spliced (top) and unspliced (bottom) isoforms. Arc color indicates probability. Black lines indicate the six common domains. Black dashed line represents the location of the intron in the unspliced isoform.

(C) Minimum free energy secondary structure model of unspliced *MUNC* color coded by SHAPE reactivities. Domains are highlighted in grey.

(D) Nucleotide-resolution SHAPE reactivity profiles for the unspliced *MUNC* isoform in-cell (top) and cell-free (bottom). Mean reactivities (± SE) are colored by relative value.

*Supplemental Figure 4*. Overexpression of Spliced Isoform of *MUNC* Leads to Induction of Promyogenic mRNAs During Differentiation

(A) Secondary structure model of spliced *MUNC* color coded for SHAPE reactivities as shown in Figure 2B.

(B-C) RT-qPCR analyses of *MUNC* constructs in B) proliferating and C) differentiating C2C12 cells. Control (Ctrl) cells do not overexpress a *MUNC* construct. Levels were normalized to *Gapdh* and are shown relative to Ctrl. Values represent at least two independent transfectants with at least three biological replicates for each of them and are presented as means ± SEM. Statistical significance was calculated using Student's t test. *** indicates $p < 0.001$ in comparison to Ctrl.

(D-F) Levels of D) *Myod1*, E) *Myog*, and F) *Myh3* in Ctrl cells and cells that express *MUNC* constructs in differentiating conditions. Levels of mRNAs were normalized to *Gapdh* and are shown relative to the wild-type spliced isoform (MS). Values represent at least two independent transfectants with at least three biological replicates for each of them and are presented as means ± SEM. Statistical significance was calculated using Student's t test. ***, **, * indicate $p < 0.001$, $<0.01$, and $< 0.05$, respectively, in comparison to MS.

(G) RT-qPCR analyses of *MUNC* constructs in differentiating cells. Levels were normalized to *Gapdh* and are shown relative to Ctrl. Values represent three biological replicates and are presented as means ± SEM. Statistical significance was calculated using Student's t test. *** indicates $p < 0.001$ in comparison to Ctrl.

(H-J) RT-qPCR analyses of levels of H) *Myod1*, I) *Myog*, and J) *Myh3* mRNAs in cells that express both ΔCH1 and ΔCH4 under differentiating conditions. Levels were normalized to *Gapdh* and are shown relative to MS. Values represent three biological replicates and are presented as means ± SEM. Statistical significance was calculated Student's t test. *** indicates < 0.001 in comparison to MS.

*Supplemental Figure 5*. Overexpression of Spliced *MUNC* Isoform Leads to Induction of Promyogenic Proteins During Differentiation

(A) Representative Western blot used for MYOD1 in differentiating control cells (Ctrl) and cells overexpressing the wild-type spliced MUNC (MS) or mutants overexpressing cells. HSP90 served as a loading control.

(B) Quantification of MYOD1 protein in differentiating cells that express indicated *MUNC* constructs normalized to HSP90 and shown relative to MS. Values represent two independent transfectants with two biological replicates for each of them and are presented as means ± SEM. Statistical significance was calculated using Student's t test. *** and * indicates $p < 0.001$ and $< 0.05$, respectively, in comparison to MS.

*Supplemental Figure 6*. Analysis of *MUNC* ChIRP-seq Datasets Reveals *MUNC* Functional Targets

(A) Venn diagram representing overlap between the two *MUNC* ChIRP datasets.

(B-C) Graphs of locations of sites of *MUNC* association in absolute distance to transcription start sites (TSS) for data generated A) for this paper and B) by Tsai et al.

(D) Workflow of analysis determining *MUNC* targets important for myogenesis.

(E) Heatmap showing clustering of data from *MUNC*-depleted cells grown under differentiation conditions (siMUNC_DM3), control cells grown under proliferating conditions (siCtrl_GM), and *MUNC*-depleted cells grown under proliferating conditions (siMUNC_GM). Microarray data from Mueller et al. (2015).

(F) Heatmap showing clustering of RNA-seq data from samples that overexpress spliced (MS_GM) and unspliced (MU_GM) isoforms grown under proliferating conditions, from samples that overexpress spliced (MS_DM3) and unspliced (MU_DM3) isoforms grown under differentiating conditions compared with data from control differentiating cells (Ctrl_DM3).

*Supplemental Figure 7*. Distinct Structural Domains of *MUNC* Regulate Expression of Different Genes.

(A-D, I-L, Q-S) ChIRP-qPCR results for 11 analyzed *MUNC* ChIRP peaks for proliferating control cells (Ctrl), *MUNC* spliced wild-type overexpressing cells (MS) and *MUNC* spliced mutants overexpressing cells (ΔCH1-L1mut). Levels were normalized to unspecific site (*Gapdh*) and shown relative to MS. Values represent two independent transfectants, three biological replicates for each of them, and are presented as means ± SEM. Statistical significance was calculated using Student's t test. ***, **, * indicates p-value < 0.001, <0.01 and < 0.05, respectively, in comparison to MS.

(E-H, M-P) RT-qPCR results for expression of 8 analyzed genes adjoining *MUNC* ChIRP peaks for proliferating control cells (Ctrl), *MUNC* spliced wild-type overexpressing cells (MS) and *MUNC* spliced mutants overexpressing cells (ΔCH1-L1mut). Levels of mRNAs were normalized to *Gapdh* and shown relative to MS. Values represent two independent transfectants, three biological replicates for each of them, and are presented as means ± SEM. Statistical significance was calculated using Student's t test. ***, **, * indicates p-value < 0.001, <0.01 and < 0.05, respectively, in comparison to MS.
